# Premorbid Characteristics of the SAPAP3-Mouse Model of Obsessive-Compulsive Disorder: Behavior, Neuroplasticity, and Psilocybin Treatment

**DOI:** 10.1101/2024.09.22.614317

**Authors:** Michal Lazar, Michal Brownstien, Alexander Botvinnik, Chloe Shevakh, Orr Shahar, Tzuri Lifschytz, Bernard Lerer

## Abstract

**Background:** SAPAP3-knockout (KO) mice develop excessive self-grooming behavior at 4-6 months of age, serving as a model for obsessive-compulsive disorder (OCD). Given that anxiety often precedes OCD diagnosis in humans, this study investigated whether juvenile SAPAP3-KO mice exhibit anxiety-like behaviors before developing the self-grooming phenotype, and whether such behaviors respond to psilocybin treatment. The study also examined four key neuroplasticity-related synaptic proteins—GAP43, PSD95, synaptophysin, and SV2A — as SAPAP3 is a postsynaptic scaffold protein that interacts with PSD95 and may affect synaptic function.

**Methods:** Two studies were conducted using male and female juvenile (10-13 weeks) SAPAP3- KO mice. Study 1 compared behavioral phenotypes between homozygous (HOM), heterozygous (HET), and wild-type (WT) mice. Study 2 evaluated a different sample of HOM and WT mice and assessed the effect of psilocybin (4.4 mg/kg) on identified behavioral differences. Both studies included comprehensive behavioral testing focused on anxiety, social interaction, and cognitive function. Additionally, levels of four synaptic proteins were measured by western blots in the frontal cortex, hippocampus, amygdala, and striatum of juvenile and adult SAPAP3-KO mice.

**Results:** In both studies, juvenile HOM SAPAP3-KO mice showed significant anxiety-like behaviors compared to WT mice, spending less time in open field center, and elevated plus maze open arms. They also buried fewer marbles and found fewer buried Oreos than WT mice. Psilocybin treatment did not improve these behavioral manifestations. Analysis of synaptic proteins revealed significant increases in GAP43, synaptophysin, and SV2A across multiple brain regions in adult male HOM mice and of SV2A in the frontal cortex of HOM females compared to WT, but not in juvenile mice of either sex.

**Conclusions:** Juvenile SAPAP3-KO mice exhibit anxiety-like behaviors before developing the characteristic excessive self-grooming phenotype, paralleling the prodromal anxiety often seen in human OCD. Unlike in adult SAPAP3-KO mice, these early manifestations were not responsive to psilocybin treatment. The age-dependent increases in synaptic proteins observed in adult but not juvenile male SAPAP3-KO mice and to a lesser extent in females, may represent compensatory plasticity changes in response to the phenotype. These results provide insights into the developmental trajectory of OCD-like behaviors and associated neuroplastic adaptations.

**Significance Statement:** Obsessive-compulsive disorder (OCD) frequently emerges during adolescence, with anxiety as a common prodromal symptom. This study investigated behavioral and molecular characteristics of juvenile SAPAP3 knockout (KO) mice, an established preclinical model of OCD, prior to manifestation of their characteristic excessive grooming phenotype. Juvenile SAPAP3 KO mice exhibited significant anxiety-like behaviors on multiple behavioral measures. While psilocybin treatment reduces OCD-like behaviors in adult SAPAP3 knockout mice, it did not ameliorate anxiety-like behaviors in juvenile mice, indicating age-dependent therapeutic effects. Notably, adult male SAPAP3 KO mice showed elevated levels of synaptic plasticity-related proteins in emotion-regulatory brain regions, whereas juvenile KO showed no such alterations. These findings demonstrate that anxiety precedes compulsive behaviors in this model and reveal age-dependent neuroplasticity changes. This developmental trajectory parallels clinical observations in OCD and provides a framework for investigating early intervention strategies.

## Introduction

Mice that carry a homozygous deletion of the SAPAP3 gene manifest, from the age of 4-6 months, a characteristic phenotype consisting of repetitive bouts of self-grooming, head-body twitches, and anxiety-related behaviors (Welch et al., 2007; Lamothe et al., 2023; Brownstien et al., 2024). The SAPAP3 knockout mouse (SAPAP3-KO) is regarded by many investigators as a relevant preclinical model of obsessive-compulsive disorder (OCD) because of the resemblance of compulsive self-grooming to compulsive behaviors manifested by patients with OCD and OCD-related phenotypes such as trichotillomania and skin picking. The model thus has reasonable face validity (defined as the similarity of what is observed in the animal model to what is observed in the human modeled organism) (Willner, 1986). The self-grooming behavior and anxiety of SAPAP3-KO mice are reduced by sub-chronic administration of the selective serotonin reuptake inhibitor (SSRI), fluoxetine (Welch et al., 2007), which is used to treat OCD, reflecting a degree of predictive validity (the extent to which the performance of the animal model in response to a defined experimental manipulation correlates with or can predict the response of the human condition to that same independent variable (Willner, 1986). Moreover, there is evidence that suggests impaired cortico-striatal connectivity in SAPAP3-KO mice, which was alleviated by ketamine administration (Davis et al., 2021). In patients with OCD, reduction of cortico-striatal hyperconnectivity was reported in patients who responded to treatment with dorsomedial prefrontal repetitive transcranial magnetic stimulation (rTMS) (Dunlop et al., 2016), suggesting that the SAPAP3-KO model of OCD may also have construct validity (a similarity in the biological underpinnings of the disease and the model) (Willner, 1986). Although not definitively validated (Belzung and Lemoine, 2011) the SAPAP3-KO model has increasing support as a model for OCD.

SAPAP3 KO mice do not manifest an excessive self-grooming phenotype before the age of 4-6 months (Welch et al., 2007; Lamothe et al., 2023). A key question is whether there are characteristic phenotypic manifestations in SAPAP3-KO mice prior to the emergence of the adult phenotype. Few studies have addressed this question directly. (Tesdahl et al., 2017) recorded ultrasonic vocalizations from 5-day-old SAPAP3—KO mice and found an increase in the number and duration of these vocalizations compared to wild-type mice. Assessments of other behavioral manifestations in SAPAP5-KO mice under 4 months old are currently lacking. Thus, it is not clear whether anxiety-like manifestations observed in adult SAPAP3 KO mice are present before the emergence of the full phenotype. This is an important question because it is well-established that there is a significant comorbidity between anxiety and OCD (Goodwin, 2015). In this context, a recent study of 206 children and adolescents showed that generalized anxiety disorder was a significant predictor of obsessive-compulsive symptoms as well as risk for OCD (Rueppel et al., 2024). Early diagnosis of OCD is important because of the possibility of early intervention, which is a focus of considerable interest (Fontenelle et al., 2022). Thus, phenotypic manifestations in SAPAP3 KO mice under 4 months old are a key unexplored question.

From the perspective of treatment, other than fluoxetine (Welch et al., 2007), there are reports that excessive self-grooming in SAPAP3-KO mice is reduced by a single administration of ketamine, the effect lasting for 3 days (Davis et al., 2021). The atypical antipsychotic, aripiprazole, was found to reduce head-body twitches and short grooming bouts in SAPAP3-KO mice but not long grooming bouts (Lamothe et al., 2023). Finally, it was recently demonstrated by (Brownstien et al., 2024) that a single administration of psilocybin or psilocybin-containing psychedelic mushroom extract significantly reduced excessive self-grooming and also head-body twitches and anxiety in male and female SAPAP3-KO mice >6 months old. The effect on self-grooming persisted for up to 42 days following a single administration.

In the current study, we examined male and female SAPAP3-KO mice aged 10-13 weeks for anxiety-related features and compared them to mice heterozygous for the mutation and wild type mice. We then examined the effect of treatment with a single dose of psilocybin on anxiety-related measures in a second cohort of juvenile SAPAP3 KO mice. In addition, we examined the levels of four proteins that play a key role in synaptic plasticity in four different brain areas in juvenile (≤3 months) and adult (>6 months) SAPAP3-KO mice. Our findings indicate that SAPAP3-KO mice manifest significant anxiety-related behavioral features before the emergence of the excessive self-grooming phenotype, but these manifestations are not responsive to treatment with psilocybin. Significant increases in brain levels of neuroplasticity-related synaptic proteins were observed in adult but not juvenile SAPAP3 -KO mice.

## Material and Methods

### Animals and Experimental Design

As previously described (Brownstien et al., 2024), a SAPAP3-KO breeding colony was established using 5 heterozygous SAPAP3-KO male and female mice kindly provided by Dr. Guoping Feng (Massachusetts Institute of Technology). Mice were housed up to 8 per cage (cage size: 43x27x30 cm) under standardized conditions with a 12-h light/dark cycle, stable temperature (22 ± 1 °C), controlled humidity (55 ± 10%), and free access to mouse colony chow and water. The cages included all genotypes under study. Mice from the same cage received the same treatment. Throughout the experiment, male and female SAPAP3-KO mice were assessed separately in order to identify any significant difference in the behavioral phenotype between the sexes. Experiments were approved by the Authority for Biological and Biomedical Models, Hebrew University of Jerusalem, Israel (Animal Care and Use Committee Approval Number: MD-21-16596-4). All efforts are made to minimize animal suffering and the number of animals used.

The purpose of Study 1 was to evaluate anxiety and depression-like phenotypes, cognition, social interaction, and social dominance in SAPAP3 HOM, HET, and WT KO mice. 44 mice homozygous (HOM) for SAPAP3-KO were used for this study (20 male, 24 female), 42 heterozygous (HET) mice (21 male, 21 female), and 55 wild-type (WT) mice (26 male, 29 female). Mice were aged 11.03±0.84 (mean ± SD) weeks at the time of evaluation (Supplementary Table 1). No treatments were administered to the mice in this study.

The purpose of Study 2, was to examine the effect of treatment with psilocybin 4.4 mg/kg i.p. on the performance of juvenile SAPAP3-KO mice on a series of tests that had shown significant differences between SAPAP3-KO genotypes in Study 1.For Study 2, 32 mice HOM for SAPAP3-KO were used (16 male, 16 female) and 32 WT mice (16 male, 16 female). HET mice were not used because of the focus on treatment effects requiring clearly differentiated groups. Mice were 11.85±0.87 (mean ±S D) weeks at the time of entry into the study and treatment administration (Supplementary Table 1). Drugs were administered by intraperitoneal (i.p.) injection in a standard injection volume in the ratio of 10 μL/1g 48 hours before behavioral tests were initiated. Mice were randomly divided into the following treatment groups: Vehicle (VEH): 0.9% saline (Male, n=16; Female, N=16); Psilocybin (PSIL): 4.4 mg/kg dissolved in the saline vehicle (Male, n=16; Female, N=16). PSIL was supplied by Usona Institute, (Madison WI, USA) and was determined by AUC at 269.00 nm (UPLC) to contain 98.75 wt. % psilocybin. The dose of psilocybin that we used (4.4 mg/kg) was chosen based on our previous dose response study on the effect of psilocybin on the head twitch response (Shahar et al., 2022; Shahar et al., 2024), and in view of the fact this dose is equivalent in mice to a 25 mg dose in humans according to the widely used DoseCal dose conversion methodology (Janhavi et al., 2022).

### Genotyping

Following the “HOTSHOT” method, genotype was determined by polymerase chain reaction (PCR) of mouse tail DNA or by mouse ear hole DNA. Three different SAPAP3 KO primers are used to distinguish the genotypes of wild type, heterozygous and Homozygous mice as follow: Primer F1 (5’ ATTGGTAGGCAATACCAACAGG 3’) and Primer R1 (5’GCAAAGGCTCTTCATATTGTTGG 3’) identify wild-type allele (around 147 base pairs), while Primer F1 and TK F2 (5’ CTTTGTGGTTCTAAGTACTGTGG 3’) identify KO allele (around 222 base pairs) (Welch et al., 2007).

### Behavioral tests

For Study 1, behavioral tests were performed according to the schedule shown in Fig. 1a. Since all the tests have objective outcome measures that are not dependent on subjective assessment, the experimenter was aware of the genotypes of the mice. The same experimenter (MB) performed all the behavioral assessments. A detailed description of the tests is provided in the Supplementary Information section of this paper.

**Fig. 1:**
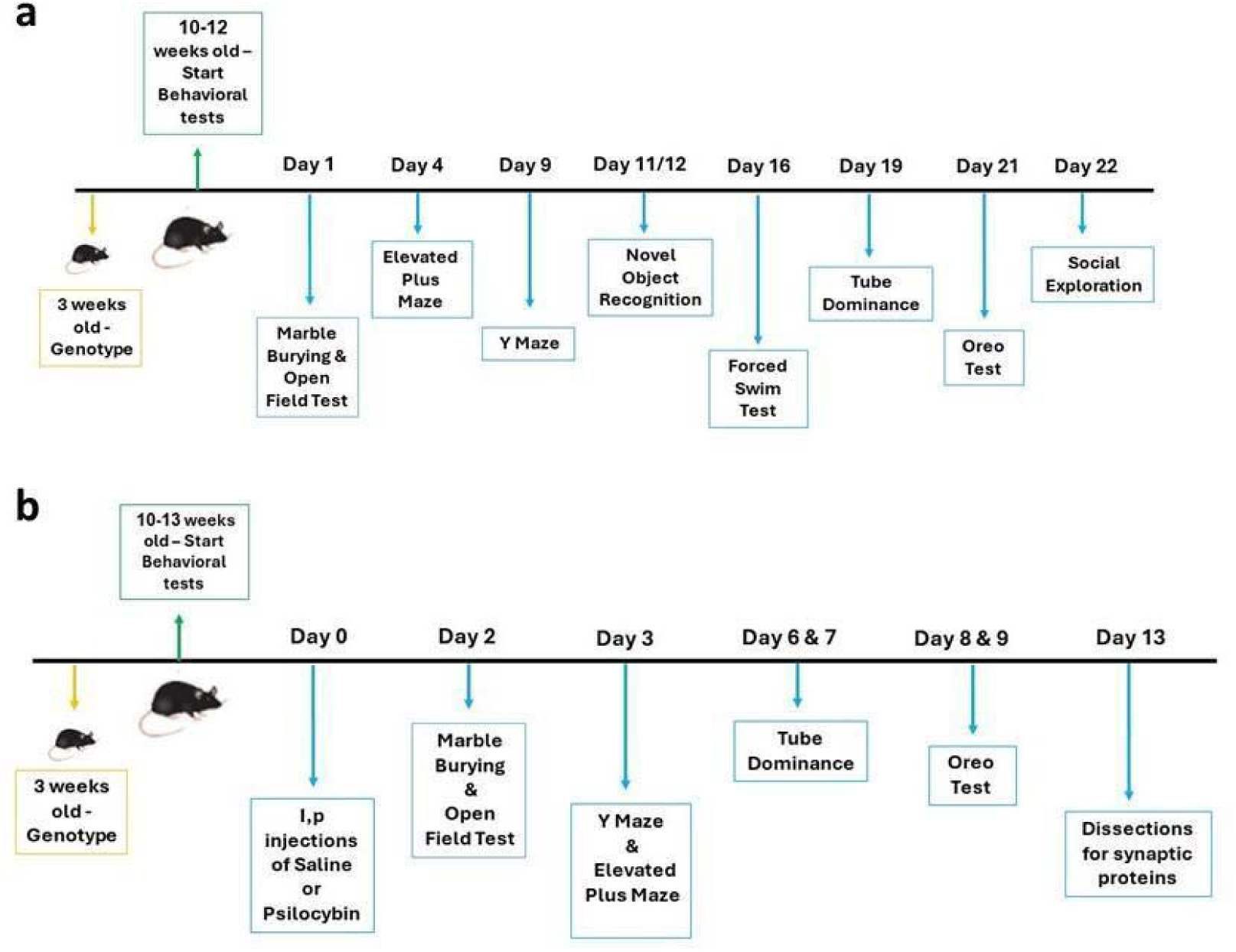
a. Timeline of the behavioral test in Study 1. **b:** Timeline of treatment and behavioral tests in Study 2.

For Study 2 behavioral tests started 48 hours after drug administration, according to the schedule shown in Fig. 1b. As in Study 1, the outcome measures were objective, and the experimenter was not blind to genotype or treatment assignment. ML performed all the assessments with the assistance of BO. The methodology applied in the tests was the same as for Study 1, as described in Supplementary Information.

### Synaptic protein assays

#### Study 1

Brain tissue samples (frontal cortex, hippocampus, amygdala, and striatum) were obtained, as previously described (Shahar et al., 2024), from 60 adult SAPAP3-KO mice aged 7 months and 16 days ± 7.8 days (males) and 7 months and 26 days ± 7.5 days (females) (mean + SD) (HOM: male n=15, female n=15; WT male n=15, female n=15;) that had not undergone behavioral testing. Hippocampus and striatum have distinct and easily identifiable morphology and were dissected fully. Frontal cortex included all cortex brain matter sectioned at coronal bregma 0.26 – 2.1 mm, and 0 – 3.5mm dorsal to ventral by brain atlas (Keith B. J. Franklin and Paxinos, 2008). Amygdala included the brain matter sectioned at coronal bregma -1.22 - -2.18 mm, 4.8 – 6mm dorsal to ventral, and 1.5 – 2.75mm medial to lateral by brain atlas (Keith B. J. Franklin and Paxinos, 2008). Samples were stored at −80 °C until assayed. Using Western blot analysis, we quantified 4 synaptic proteins (GAP43, PSD95, synaptophysin, and SV2A) in each of the 4 brain areas, as described in our previous report (Shahar et al., 2024). Briefly, the brain tissue samples were lysed in Pierce RIPA sample buffer (Thermo Scientific, USA), supplemented with protease inhibitor cocktail (Roche Diagnostics, Germany) and boiled for 10 min. Equivalent amounts of protein extracts (20 mg) were analyzed by SDS–12% PAGE, followed by transfer of the proteins to polyvinylidene fluoride membrane. Blots were blocked in 5% fat free milk in TBST buffer (Tris-Tween-buffered saline) and incubated in primary antibodies, one hour at room temperature. Primary antibodies included rabbit anti-GAP43 (ab75810, 1:2000; Abcam, UK), rabbit anti-PSD95 (ab238135, 1:2000; Abcam, UK), rabbit anti-Synaptophysin (ab32127, 1:2000; Abcam, UK), rabbit anti-SV2A (ab54351, 1:100; Abcam, UK) and mouse anti-β-Actin (8H10D10, 1:5000, Cell Signaling Technology). Blots were washed 3 times and incubated with horseradish peroxidase-conjugated secondary antibodies (1:5000, ABclonal, China) for 1 h, followed by repeated washing with TBST buffer. Proteins were visualized by using enhanced chemiluminescence (ChemiDoc Reader MP, Bio-Rad, USA). The amount of each phosphorylated protein was normalized to the amount of the corresponding total protein detected in the sample and measured by intensity of β-actin. (Representative immunoblots are shown in Supplementary Figure 14A-B)

#### Study 2

On day 13 after drug administration, brain tissue samples (frontal cortex, hippocampus, amygdala and striatum) were obtained from 24 juvenile SAPAP3-KO mice aged 12.96±0.999 (mean ± SD) weeks (HOM: male n=5, female n=6; WT male n=7, female n=6;) SAPAP3-KO (n=11) and WT mice (n=13) that had been administered VEH, and were stored at −80 °C. Synaptic proteins (GAP43, PSD95, synaptophysin and SV2A) were quantified as described above. (Representative immunoblots are shown in Supplementary Figure 14C- D).

### Statistical analysis

The experimental data are presented in all figures as the meanlll±lllstandard error of the mean (SEM) and in the text as the meanlll±lllstandard deviation (SD). To determine inter-group differences, student t-tests, or one or two- or three-way analysis of variance (ANOVA) were used as indicated. In Study 1 Sidak tests were used to analyze post-hoc comparisons. plll<lll0.05 (two-tailed) was the criterion for significance. Male and female mice were included in both studies (Beery and Zucker, 2011) and results were tested for differences between the sexes by two (Study 1) or three way ANOVA (Study 2). Correlations were tested by the Pearson Product Moment Correlation Test. Graph Pad Prism, version 9.3.1 software was used for all statistical analyses. A nested t-test was performed to compare the expression levels of all 4 synaptic proteins (GAP43, PSD95, synaptophysin, SV2A) in each brain area (frontal cortex, hippocampus, amygdala and striatum) between two genotypes (SAPAP3 KO and WT) and of each synaptic protein over all 4 brain areas. Statistical significance was assessed using a t-test (two-tailed). p < 0.05 was considered statistically significant. Results are shown as mean ± SEM.

## Results

### Behavioral Tests

#### Study 1

In Study 1, SAPAP3 HOM, HET, and WT KO mice underwent a series of behavioral tests to evaluate anxiety and depression-like phenotypes, cognition, social interaction, and social dominance. The schedule for these tests is given in Fig 1a. The results of this study show significant effects of the SAPAP3-KO genotype on several behavioral tests.

We focused on two key measures yielded by the open field test (OFT) – total distance travelled in the 10 minutes of the test and total time spent in the central area of the arena. On the OFT (Fig. 2-Ia. Fig. 2-Ib), two-way ANOVA showed a significant effect of genotype on total distance travelled (F=20.11; df=2, 119, p<0.0001) but no significant effect of sex. Further analyses were conducted in males and females separately. One-way ANOVA of total distance traveled showed a significant effect of genotype in both males (F=7.89, df=2,63, p=0.0009) and females (F=12.71, df=2,56, p<0.0001). Sidak post-hoc tests showed that male HOM SAPAP3-KO mice traveled a significantly lower distance (p=0.0007) than male WT mice (Fig2-Ia), and female HOM SAPAP3- KO mice traveled a significantly lower distance than female WT (p=0.0007) and HET mice (p=0.04) (Fig2-Ib).

**Fig. 2:**
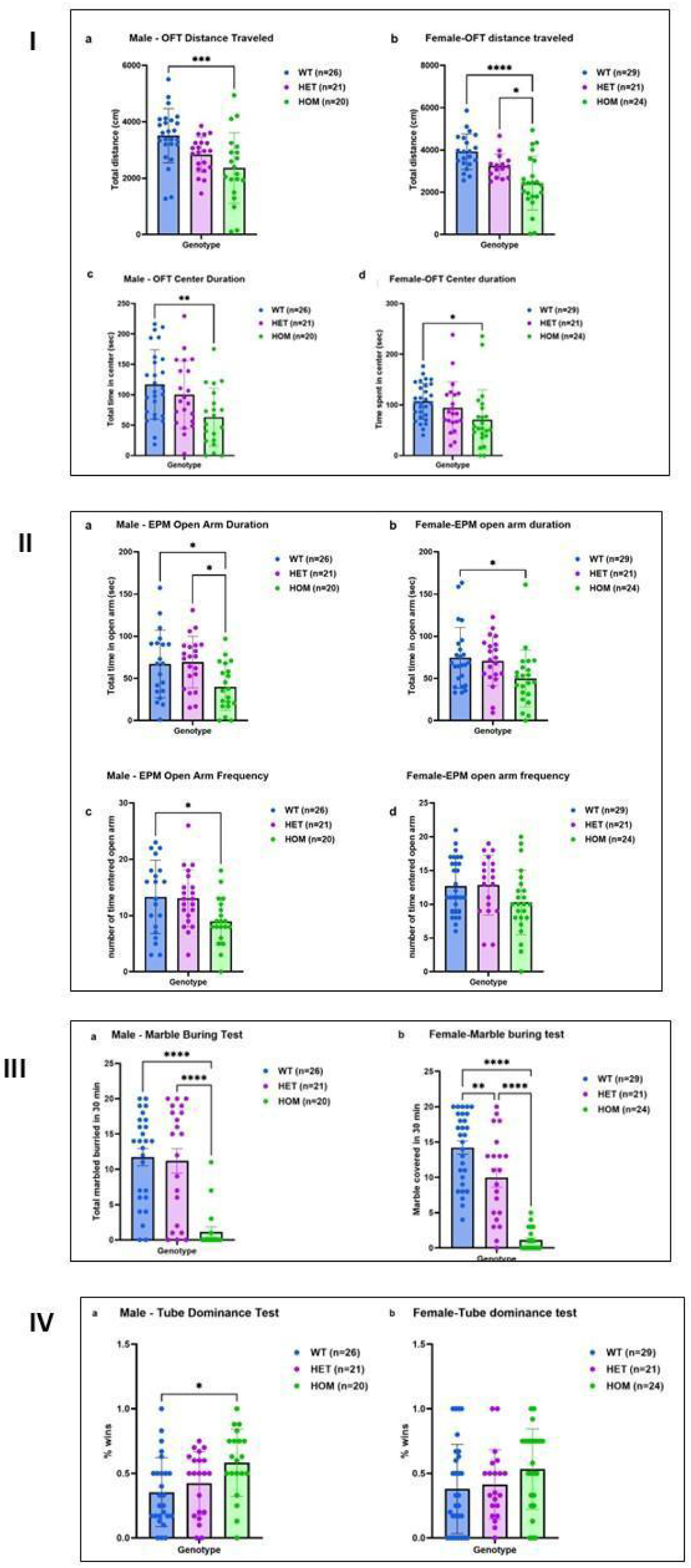
I. Open field Test comparing WT, HET, and HOM SAPAP3-KO mice: **a, b** - Distance traveled by males (a) and females (b). Significant effect of genotype in a) males (F=7.89; df 2,63, p=0.0009) and b) females (F=12.71; df 2,56, p<0.0001). **c, d.** - Center duration of males (c) and females (d) Significant effect of genotype in c) males (F=5.749, 2,64, p=0.0051) and d) females (F=3.687; df 2,70, p=0.030). Sidak post-hoc test: *p<0.05, **p<.01, ***p<.001,****p<.0001 II. Elevated Plus Maze comparing WT, HET and HOM SAPAP3-KO mice: **a, b** – Time spent in open arms in males (a) and females (b). Significant effect of genotype in a) males (F=4.874, df 2,58 p=0.001) and b) females (F=3.556, df 2,54, p=03). **c, d**. – Number of times entered open arms in males (c) and females (d). Significant effect of genotype in c) males (F=4.19, df 2,58, p=0.02) Sidak post hoc tests: *p<.05, **p<.02, ***p<.001 III. Marble burying test in males (a) and females (b), comparing WT, HET, and HOM SAPAP3-KO mice. Significant effect of genotype in a) males (F=18.90; df 2,63, p<0.0001) and b) females (F=53.74; df 2,71, p<0.0001). Sidak post-hoc test: **p<.01, ****p<.0001 IV. Tube Dominance test in males (a) and females (b), comparing WT, HET and HOM SAPAP3- KO mice percentage of wins. Significant effect of genotype in a) males (F=4.516, df 2,64, p=0.01) Sidak post hoc test: *p<.05

For time spent in the center of the open field, two-way ANOVA showed a significant effect of genotype on total distance traveled (F=9.54; df=2, 134, p=0.0001) but no significant effect of sex. One-way ANOVAs showed a significant effect of genotype on time spent in the center of the open field for male (F=5.74, df=2,64, p=0.005) and female (F=3.687; df=2,70; p=0.03) SAPAP3-KO mice (Fig. 2-Ic, Fig. 2-1d). Male (p=0.004) and female (p=0.026) HOM SAPAP3-KO mice both spent significantly less time in the center of the OFT arena than their WT littermates (Fig. 2-Ic, Fig. 2-Id), indicating a greater level of anxiety. Regarding time spent in the periphery of the open field, one-way ANOVA showed a significant effect of genotype in male mice (F=7.46, df=2,59, p=0.0013). Male HOM SAPAP3-KO mice spent significantly more time in the periphery (p=0.0009) than male WT mice. For the female group, there were no significant differences in time spent in the periphery.

On the elevated plus maze (EPM), time spent in the unprotected open arms and frequency of entry into these arms are indicative of level of anxiety, with more anxious mice spending less time in these arms and entering less frequently. Two-way ANOVA showed a significant effect of genotype but not sex on time spent in the center of the open field (F=8.25; df=2, 122, p=0.0004). One way ANOVA showed a significant effect of genotype on time spent in the open arms of the EPM for male (F=4.874, df=2,58, p=0.001) and female (F=3.55, df=2,64, p=0.034) SAPAP3-KO mice (Fig. 2-IIa, Fig. 2-IIb). Sidak post hoc tests showed that male HOM SAPAP3-KO mice spent significantly less time in the open arms than HET SAPAP3-KO (p=0.03) and WT mice (p=0.01) (Fig. 2-IIa) and also female HOM SAPAP3-KO mice compared to female WT mice (p=0.04) (Fig. 2-IIb). One-way ANOVA also showed a significant effect of genotype on the number of times mice entered the open arms for male SAPAP3-KO mice (F=4.19, df=2,58, p=0.02). The total number of times entered into the open arm was significantly lower in male HOM SAPAP3-KO mice (p=0.03) compared to WT.

On the marble burying (MB) test (Fig 2-III), the key measure is number of marbles buried over 30 minutes. Two-way ANOVA showed a significant effect of genotype on number of marbles buried (F=60.13; df=2, 134, p<0.0001) but no significant effect of sex. One-way ANOVA showed a significant genotype effect for both males and females (males: F=18.9, df=2,63, p<0.0001; females: F=53.74, df=2,71, p<0.0001). Sidak post-hoc tests showed that male and female HOM SAPAP3-KO mice buried significantly fewer marbles than their WT littermates (p<0.0001) and female HOM significantly fewer than female HET mice (p<0.01) (Fig. 2-IIIa and b). To determine whether lower marble burying in the SAPAP3 KO mice could be related to the activity level of the mice, we examined the correlation between the number of marbles buried and the distance covered in the OFT. The correlation was significant (r=0.35, p<0.0001, 140 XY pairs), indicating that mice that are less active bury fewer marbles and vice versa. We further examined whether marble burying could be related to the level of anxiety of the SAPAP3 KO mice as reflected in the time spent in the open arms of the EPM. This correlation was also significant (r=0.22, p=0.007, 140 XY pairs) indicating that mice that are more anxious bury fewer marbles. There was a trend for distance covered in the OFT and time spent in the open arms of the EPM to be related, but this was not significant (r=0.27, p=0.07, 140 XY pairs).

Performance on the tube dominance test reflects social assertiveness as reflected in percent of winning (Zhou et al., 2017). Two-way ANOVA showed a borderline significant effect of genotype on winning (F=5.48; df=2, 136, p=0.05) but no significant effect of sex and no significant genotype x sex interaction. There were no significant differences between genotypes in the female group (Fig. 2-IVb). In male mice, one-way ANOVA showed a significant effect of genotype (F=4.516, df=2,64; p=0.01) with a higher percent of winning (p=0.01) in HOM compared to WT littermates. (Fig. 2- IVa). Percent of winning was not related to activity level (distance traveled on the OFT) (r=- 0.03, p>0.01, 140 XY pairs) nor to level of anxiety (time spent in the open arms of the EPM) (r=0.01, p>0.01, 140 XY pairs)

The Oreo test is based on the preference of mice for sweet objects (Markov, 2022). The first assessment was whether previously habituated mice placed in a cage with a buried Oreo would find the cookie and dig it up and eat it. There was a significant effect of genotype on this parameter among male mice (Table 1). 88.96% of male WT mice found the Oreo and 81.82% of male HET mice but only 35.00% of male HOM mice (x =16.75, df 2, p=0.0002). Among female mice, the differences were not significant (WT 62.07% found; HET 61.90% found; HOM 58,33% found; x =0.10, df 2, p=0.93). There was no significant effect of genotype on Oreos eaten by those mice that found them among males or female mice (Table 1a). Number of Oreos found and number of Oreos eaten were not related to activity level (distance traveled on the OFT) (r=0.07, p>0.1, 140 XY pairs) nor to level of anxiety (time spent in the open arms of the EPM) (r=0.07, p>0.1, 140 XY pairs)

**Table 1:**
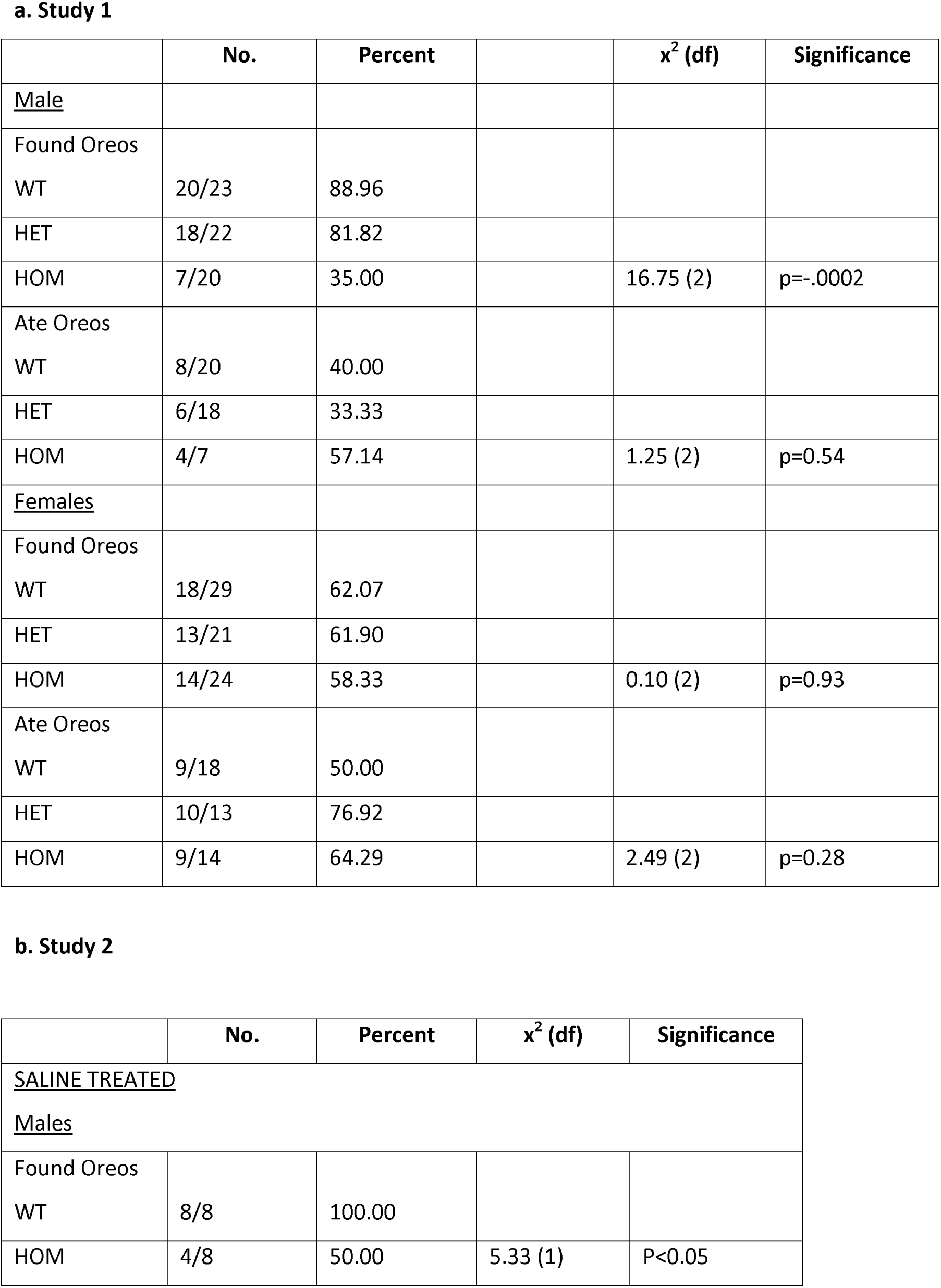
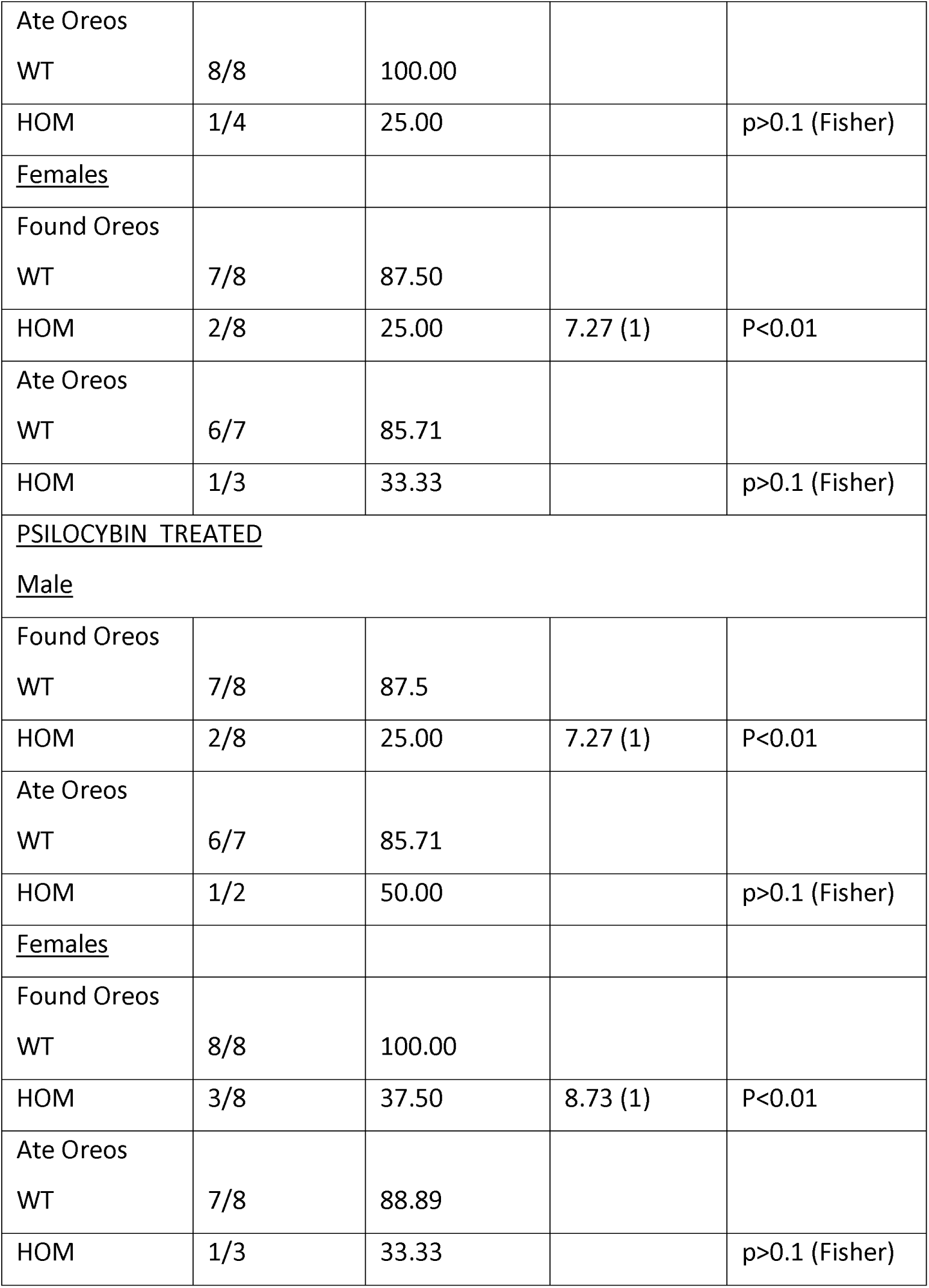
Effect of SAPAP3-KO genotype on finding and eating Oreo Cookies (No./% of mice that found cookies in the test and number of those which found cookies that ate them)

Contrary to these positive findings, other tests did not show an effect of SAPAP3-KO genotype in male or female nice. On the forced swim test (FST), there was no significant effect of genotype on duration (Supplementary Fig 1a, 1b) or frequency (Supplementary Fig. 1c, Fig. 1d) of inactivity in male or female mice. The social exploration test (Supplementary Fig. 2a, 2b), the NOR test (Supplementary Fig. 2c, 2d), and the Y maze (Supplementary Fig. 2d, 2e) also showed no significant differences between genotypes in both male and female groups.

#### Study 2

In Study 2, we examined the effect of treatment with psilocybin 4.4 mg/kg i.p. on the performance of juvenile SAPAP3-KO mice (mean age 11.85±0.86 weeks) on a series of tests that had shown significant differences between SAPAP3-KO genotypes in Study 1. The study timeline and test battery are summarized in Fig 1b.

There were no significant effects of psilocybin treatment on any of the behavioral measures, as indicated by the lack of a significant main effect of treatment on three way that took sex into account and on two-way ANOVA in the sexes separately (see below). However, there were significant effects of genotype over and above treatment and significant post-hoc effects in both saline and psilocybin-treated mice on a number of tests.

On the OFT (Fig. 2-Ia. Fig. 2-Ib), which examines activity and anxiety-like behaviors, three-way ANOVA showed a significant effect of genotype on total distance travelled (F=156.00; df=1, 56, p<0.0001) but no significant effect of sex or treatment with psilocybin. Further analyses were conducted in males and females separately. (Fig. 3-Ia and b). There was a significant effect of genotype over and above treatment on distance traveled in male (F=8.121; df 1,28; p=0.0081) (Fig. 3 I-a) and female (F=32.35; df 1,28; p<0.0001) (Fig. 3 I-b) SAPAP-3 KO mice. Tukey’s multiple comparisons test showed significantly lower distance traveled by male (p<0.05, Psilocybin WT vs. Psilocybin HOM) (Fig. 3 1-a) and female HOM vs. WT mice (p<0.01, Saline WT vs. Saline HOM; p=0.0014, Psilocybin WT vs. Psilocybin HOM) (Fig. 3 I-b) SAPAP3-KO mice. The center duration of SAPAP3 KO mice was not significantly shorter than WT mice in males (Fig. 7e) but was significantly so in females (F=9.12; df 1,28; p=0.0053; p<0.05, Psilocybin WT vs. Psilocybin HOM) (Fig. 3 I-d). For periphery duration there was no significant difference between genotypes in males or females.

**Fig. 3:**
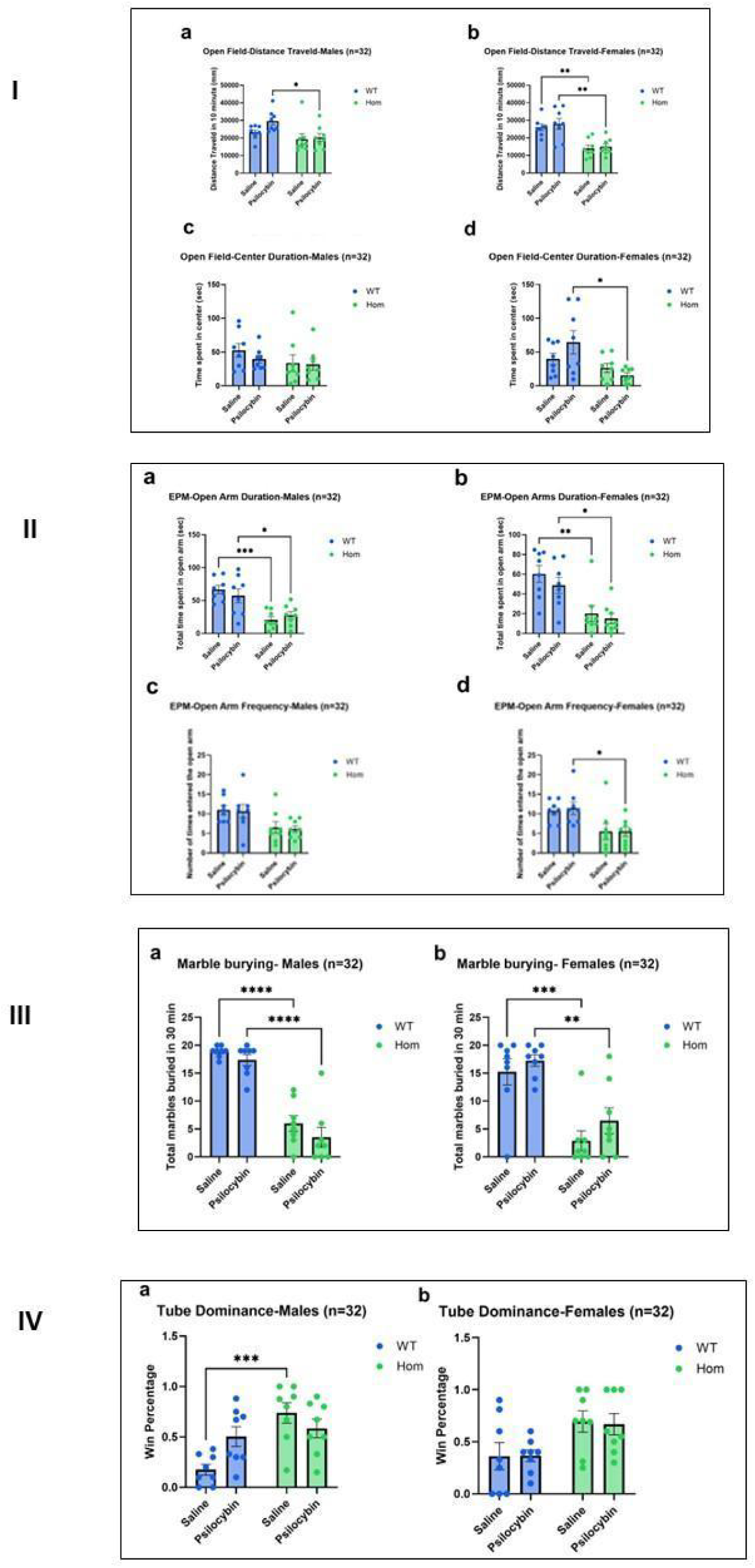
I. Open field Test comparing WT and HOM SAPAP3-KO mice with and without treatment with psilocybin: **a, b** -Distance traveled: Significant effect of genotype in a) males (F=8.121; df 1,28; p=0.0081) and b) females (F=32.35; df 1,28; p<0.0001). **c, d** -Center Duration: No significant effects in a) males. Significant effect of genotype in b) females (F=9.12; df 1,28; p=0.0053). *p<0.05, **p<0.01 (Dunnet’s test) II. Elevated Plus Maze Test comparing WT and HOM SAPAP3-KO mice with and without treatment with psilocybin: **a,b** - Open arms duration of a) Males (F=27.76; df 1,28; p<0.0001; p=0.0005), b) Females (F=23.46; df 1,28; p<0.0001). **c,d** - Open arm frequency of c) males (F=11.15; df 1,28; p=0.002); d) females (F=14.19; df 1,28; p=0.0008). *p<0.05, **p<0.01, ***p<0.001 (Dunnet’s test) III. Marbles buried in 30 minutes comparing WT and HOM SAPAP3-KO mice with and without treatment with psilocybin: **a-** Marbles buried in 30 minutes by males (F=113.6, df 1, 28, <0.0001). **b-** Marbles buried in 30 minutes by females (F=35.40, df 1, 28, p<0.0001). **p<0.01, ***p<0.001, ****p<0.0001 (Dunnet’s test) IV. Tube Dominance Test in males (a) and females (b) in WT and SAPAP3 KO mice, with our without treatment with psilocybin: **a.** - Males (F=13.28; df 1,28; p=0.001) **b.** - Females (F=9.86; df 1,28; p=0.003) ****p<0.0001 (Dunnet’s test)

On the EPM three-way ANOVA showed a significant effect of genotype on total distance travelled (F=50.97; df=1, 56, p<0.0001) but no significant effect of sex or treatment. Further analyses were conducted in males and females separately. R esults (Fig. 3 II) revealed a striking level of anxiety among HOM SAPAP3-KO mice of both sexes compared to WT mice, as reflected by the effect of genotype on time spent in the open arms of the apparatus (males: F =27.76, df 1,28, p<0.0001; females: F=23.46, df 1,28, p<0.0001). Tukey’s multiple comparisons test showed that v ehicle-treated HOM mice of both sexes spent significantly less time in the open arms of the EPM than vehicle-treated WT mice (males: p=0.0005; females: p=0.004), and also psilocybin-treated HOM mice compared to psilocybin-treated WT mice (males: p=0.03; females: p=0.02) (Fig. 3 II-a, b). For time spent in the closed arms of the EPM, there was also a significant effect of genotype in both sexes (males: F=7.74; df 1,28; p=0.01; females: F=19.17; df 1,28; p=0.0002); post hoc tests were significant only for vehicle-treated HOM female mice (p=0.048) that spent more time in the closed arms than vehicle-treated WT female mice. The effect of genotype on frequency of entrances to the open arms was significant in males (F=11.15; df 1,28; p=0.002; and females (F=14.19; df 1,28; p=0.0008; p<0.039, Psilocybin WT vs. Psilocybin HOM) (Fig. 3 II-d). The difference in frequency of entrances to closed arms was significant between genotypes in females (F=19.64; df 1,28; p=0.0001; p<0.01, Saline WT vs. Saline HOM; p<0.05, Psilocybin WT vs. Psilocybin HOM), but not in males.

As in Study 1, the marble burying test (Fig. 3-III) yielded striking results. Three-way ANOVA showed a significant effect of genotype on number of marbles buried (F=116.20; df=1, 56, p<0.0001) but no significant effect of sex or treatment. Further analyses were conducted in males and females separately by two-way ANOVA. There was a highly significant genotype effect in both sexes (males: F=113.6, df 1, 28, p<0.0001; females: F=35.40, df 1, 28, p<0.0001). Tukey post-hoc tests showed that v ehicle-treated HOM mice of both sexes buried significantly fewer marbles than WT mice (males p<0.0001; females p=0.0006), as was the case for psilocybin-treated HOM mice compared to psilocybin-treated WT mice (males: p<0.0001; females p=0.0028) (Fig. 3-IIIa, Fig. 3 III-b). As in Study 1, we sought to determine whether lower marble burying in the SAPAP3 KO mice could be related to the activity level of the mice. We examined the correlation between the number of marbles buried and the distance covered in the OFT. The correlation was significant (r=0.45, p=0.0002, 64 XY pairs); mice that covered less distance of the OFT buried fewer marbles and vice versa. We further examined whether lower marble burying could be related to the level of anxiety of the SAPAP3 KO mice as reflected in the time spent in the open arms of the EPM. This correlation was also significant (r=0.57, p=<0.0001, 54 XY pairs); mice that spent less time in the open arms of the EPM buried fewer marbles and vice versa. It is noteworthy that distance traveled on the OFT and time spent in the open arms of the EPM were correlated (r=0.52, p<0.0001, 54 XY pairs) i.e. greater activity was associated with less anxiety and vice versa.

On the Tube Dominance test (Fig. 3 IV),. Three-way ANOVA showed a significant effect of genotype on percent of winning (F=18.70; df=1,62, p<0.0001) but no significant effect of sex or treatment. Further analyses were conducted in males and females separately by two-way ANOVA. Male SAPAP3 KO mice were significantly more dominant than WT mice (F=13.28; df 1,28; p=0.001; ***p<0.001, Saline WT vs. Saline HOM) (Fig. 3. IV-a); the effect of genotype was also significant in females (F=9.86; df 1,28; p=0.003). We examined the correlation between the percentage of winning and the distance covered in the OFT. The correlation was significant but inverse (r=-0.42, p=0.0003, 70 XY pairs) i.e. mice that won more frequently covered less distance on the OFT and vice versa. We further examined whether the percentage of winning could be related to the level of anxiety of the SAPAP3 KO mice as reflected in the time spent in the open arms of the EPM. This correlation was also significant and inverse (r=-0.31, p=0.013, 63 XY pairs), indicating that mice that are less anxious (more time in the open arms of the EPM) have a higher percentage of winning and vice versa.

The Oreo test results in Study 2 (Table 1b) were similar to those in Study 1. Whether treated with saline or psilocybin, WT mice found the Oreo cookie significantly more frequently than HOM mice. The percentage of mice succeeding was >85% in all cases. For HOM mice, the percentage finding the Oreo was 50% or less. Comparisons between WT and HOM for Oreos were statistically significant for both sexes. For Oreos eaten, there was a clear numerical trend for WT mice to eat more of the Oreos found but the number of observations was too small to permit statistical significance to be demonstrated. We examined the correlation between the percentage of Oreos found and the distance covered in the OFT. The correlation was significant (r=0.41, p=0.0007, 64 XY pairs) i.e. mice that were more were active found more Oreos. We further examined whether the percentage of Oreos found could be related to the level of anxiety of the SAPAP3 KO mice as reflected in the time spent in the open arms of the EPM. This correlation was also significant (r=0.52, p<0.0001, 59 XY pairs) i.e. mice that were less anxious (more time in the open arms of the EPM) found more Oreos.

On the Y Maze which measures short-term special memory (Kitanaka et al., 2015), there were no significant effects of genotype on direct and indirect revisits (Supplementary Fig. 3a-d). 2-way ANOVA showed a significant effect of genotype on max alternations in females (F=7.77; df 1,28; p=0.009) (Fig 4f) but not males (Supplementary Fig. 3e). On Tukey’s multiple comparisons post-hoc tests, complete alternations showed significant difference between genotype in males (F=11.85, df 1,28, p=0.001; *p<0.05, Saline WT vs. Saline HOM) (Supplementary Fig. 3g) and females (F=10.09; df 1,28; p<0.003) (Supplementary Fig 3h). However, the complete alternation ratio did not show a significant difference between genotypes (Supplementary Fig. 3i).

**Fig. 4.**
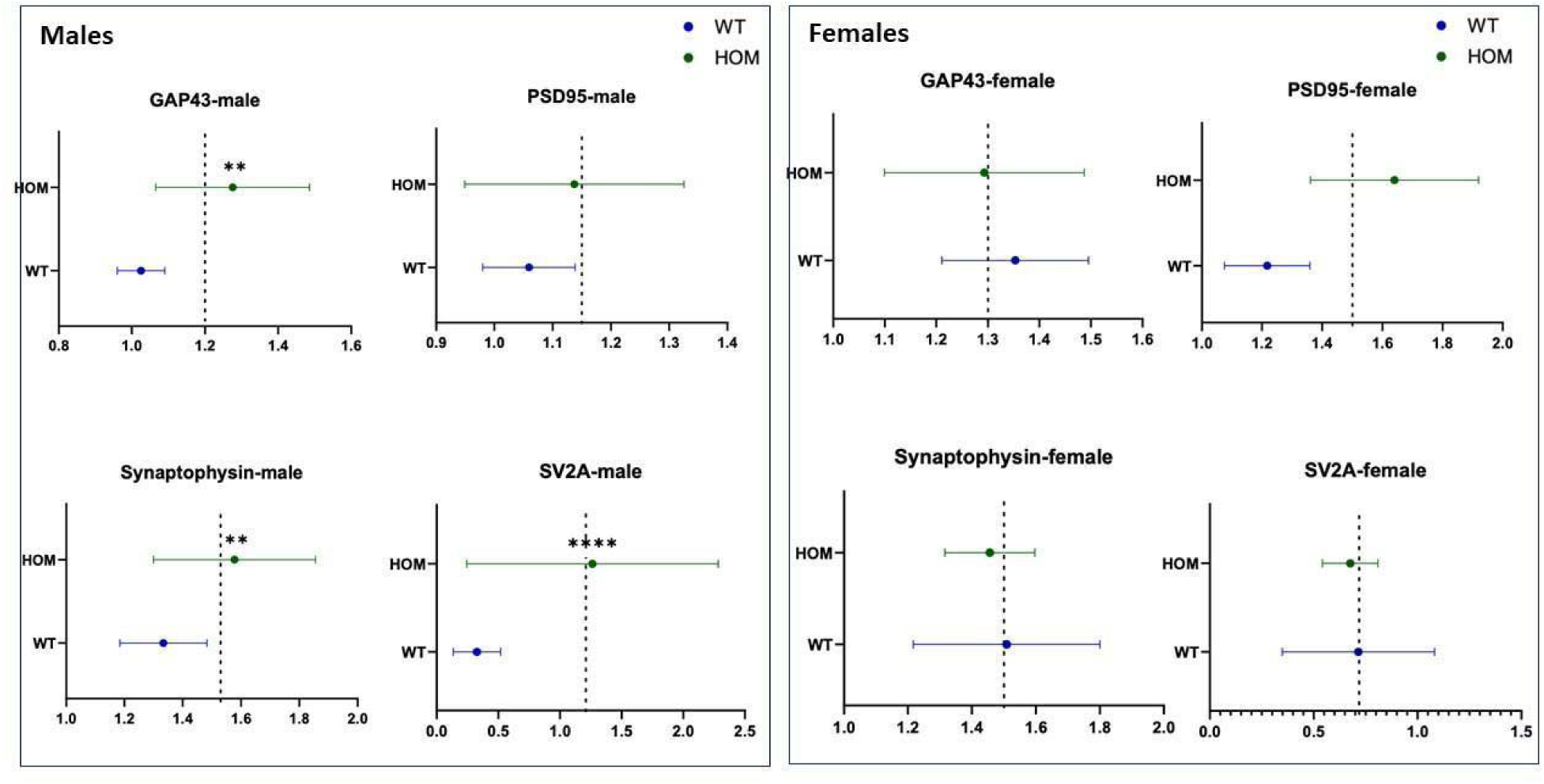
Study1: Nested analysis of synaptic proteins (GAP43, PSD95, synaptophysin, and SV2A) by genotype (SAPAP3-KO and WT) across 4 brain areas (frontal cortex, hippocampus, amygdala, and striatum) in male (left) and female (right) mice. Statistical significance was assessed using t- test (two tailed). Results are shown as mean ± SEM. **p<0.01, ****p<0.0001

### Synaptic Proteins

#### Study 1

Effects of sex and genotype on levels of 4 synaptic proteins (GAP43, PSD95, synaptophysin, and SV2A) were analyzed in the 4 brain areas we studied (frontal cortex, hippocampus, amygdala and striatum) by two-way ANOVA (See Supplementary Figures 4-7). The synaptic protein that showed the strongest effect of genotype was SV2A (Supplementary Fig. 7). Two-way ANOVA showed a significant effect of genotype on SV2A levels in the frontal cortex (F=17.22, df 1,55, p=0.001), with female HOM mice manifesting significantly higher SV2A levels than female WT mice (Tukey post-hoc, p=0.01). Genotype effects were also highly significant for SV2A in the hippocampus (F=14.67, df 1,56, p=0.0003), amygdala (F=14.48, df 1,53, p=0.0004), and striatum (F=4.59, df 1,53, p=0.03). Post-hoc tests showed that SV2A levels were significantly higher in male HOM than male WT mice in all three areas (p<0.0001). Significant effects of genotype were observed for GAP43 in the frontal cortex (F=7.30, df 1,56, p=0.009) and amygdala (F=8.24, df 1,54, p=0.005); post-hoc significance for male HOM vs. male WT was 0.02 and 0.04 for each area respectively (Supplementary Fig. 4). For PSD95 there was a significant effect of genotype in the frontal cortex (F=6.70, df 1,54, p=0.01) (Supplementary Fig. 5), and for synaptophysin in the amygdala (F=7.34, d 1,56, p=0.008), but post-hoc tests were not significant in either sex separately (Supplementary Fig. 6).

There were significant sex effects on SV2A levels in the amygdala (F=12.32, df 1,53, p=0.0009) and striatum (F=7.48, df 1,53, p=0.008) (Supplementary Fig. 7); on GAP43 in the amygdala (F=6.15, df 1,54, p=0.01) (Supplementary Fig. 4); on PSD95 in the frontal cortex (F=5.43, df 1,54, p=0.02) and hippocampus (F=4.38, df 1,56, p=0.04) (Supplementary Fig. 5); and on synaptophysin in the striatum (F=11.77, df 1,55, p=0.001) (Supplementary Fig. 6). A significant sex x genotype interaction was observed for SV2A in the hippocampus (F=20,94, df 1,56, p=,0.0001), amygdala (F=26.75, df 1,53, p<0.0001) and striatum (F=15.08, df 1,53, p=0.0003) (Supplementary Fig. 7) and for synaptophysin the frontal cortex (F=13.51 df 1,55, p=0.005) (Supplementary Fig. 6).

To analyze synaptic proteins over all four brain areas, nested t-tests were performed (Fig. 4) comparing each synaptic protein in HOM and WT mice over all brain areas. Nested t-tests were performed separately for male and female mice. In males, the analysis showed significantly higher levels of GAP43 (p=0.001), synaptophysin (p=0.003), and SV2A (p<0.0001) but not of PSD95 across the 4 brain areas in HOM vs WT SAPAP3-KO mice (Fig. 4, Left Panel). None of the differences were significant for female SAPAP3-KO mice (Fig. 5, Right Panel). Examining all 4 synaptic proteins within each brain area separately, significantly higher synaptic protein levels were observed in male HOM SAPAP3-KO mice than in WT mice in the frontal cortex (p=0.01), hippocampus (p=0.01) and amygdala (p=0.001) but not striatum (Fig. 5, Left Panel). No significant differences were observed in any of the brain areas in female mice (Fig. 5, Right Panel).

**Fig. 5.**
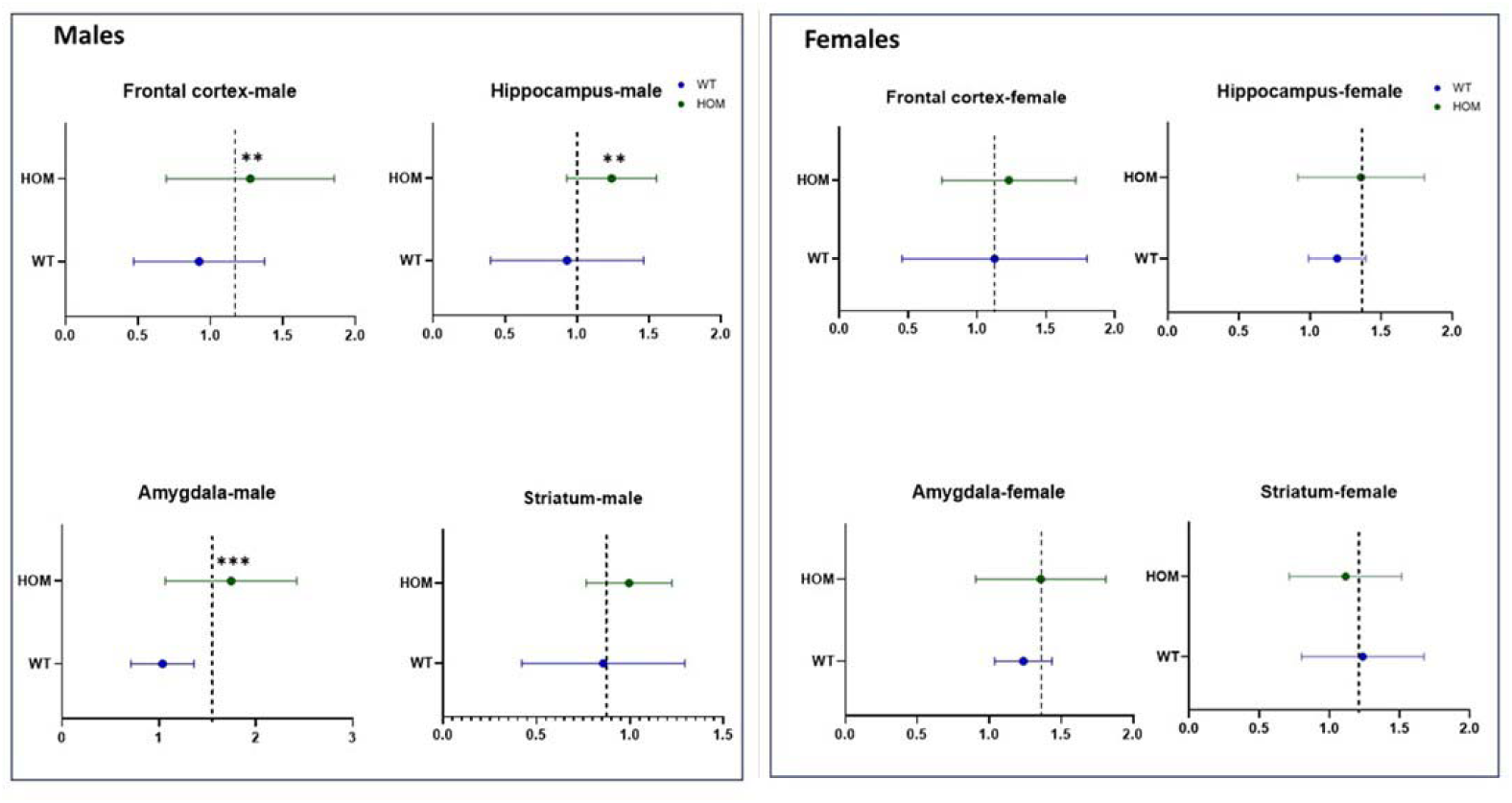
Nested analysis of 4 synaptic proteins (GAP43, PSD95, synaptophysin, and SV2A) within each of 4 brain areas (frontal cortex, hippocampus, amygdala, striatum) in male (left) and female (right) HOM vs. WT SAPAP3-KO mice. Statistical significance was assessed using t-test (two tailed). Results are shown as mean ± SEM. **p<0.01, ***p<0.001

#### Study 2

We analyzed synaptic protein neuroplasticity markers on brain samples obtained from 11 HOM (5 males and 6 females for each genotype) and 13 WT (7 males and 6 females for each genotype) vehicle-treated mice on day 13 after treatment administration. PSIL-treated mice were not examined. We performed two-way ANOVA for each of the 4 synaptic proteins in each of the 4 brain areas, with sex and genotype as factors (Supplementary Figures 8-11). This analysis revealed a significant sex effect for PSD95 in the amygdala (F=7.10, df 1,16, p=0.01), a significant sex by genotype interaction for PSD95 in the hippocampus (F=7.54, df 1,20, p=0.01) and a significant sex by genotype interaction for synaptophysin in the striatum (F=7.49, df 1,20, p=0.008). No post-hoc comparisons of HOM vs. WT genotype in males or females were significant for any of the significant sex effects or sex x genotype interactions. We performed a nested t-test comparing each synaptic protein (GAP43, PSD95, synaptophysin, SV2A – Supplementary Fig. 13) in HOM versus WT mice over all brain areas (frontal cortex, hippocampus, amygdala and striatum). This analysis did not show significant differences between genotypes for any of the synaptic proteins across all brain areas. For analysis of the synaptic protein neuroplasticity markers within each brain area, a nested t-test was performed comparing levels of all four synaptic proteins in HOM and WT mice in each brain area (frontal cortex, hippocampus, amygdala, striatum – Supplementary Fig. 14). This analysis did not show significant differences over all synaptic proteins within any of the brain areas.

## Discussion

Age of onset in OCD has been reported to be bimodal, with a mean of 12.8 (S.D. = 4.9) years in the early-onset group and 24.9 (S.D. = 9.3) years in the late-onset group (Anholt et al., 2014). Anxiety is a common comorbid feature of OCD (Tesdahl et al., 2017), and prodromal anxiety symptoms are frequent in youth (Rueppel et al., 2024) and adults (Fava et al., 1996) who subsequently develop OCD. Since the excessive self-grooming observed in SAPAP3-KO mice is regarded as a possible rodent model of OCD-like behavior, it is of interest to know whether SAPAP3-KO mice below the age at which the characteristic phenotype is observed manifest behavioral and other abnormalities. This question was the focus of the current study in which SAPAP3-KO mice of both sexes were assessed for a series of behavioral characteristics, including anxiety, depressive-like features, cognitive function, sociability, and social dominance. The effect of treatment with psilocybin on these behavioral features was evaluated. In addition, we measured levels of synaptic proteins in four brain regions in mice homozygous for the SAPAP3 deletion compared to wild-type mice as markers of neuroplasticity (Shahar et al., 2024).

Our results in Study 1 for time spent in the center of the open field and time spent in the open arms of the EPM, show clear evidence for higher levels of anxiety in in male and female homozygous SAPAP3-KO mice compared to WT mice. These findings were supported by those of Study 2, for females in regard to center duration in the open field and for both sexes in regard to time spent in the open arms of the EPM. This evidence for higher anxiety levels in juvenile SAPAP3-KO mice is in accordance with findings of (Welch et al., 2007) and (Brownstien et al., 2024) in adult SAPAP3 KO mice that manifest the full self-grooming and head-twitch phenotype. (Welch et al., 2007) found that 6 days of treatment with fluoxetine significantly alleviated anxiety like features on the elevated plus maze. (Brownstien et al., 2024) found that a single treatment with psychedelic mushroom extract (containing 4.4 mg/kg psilocybin) significantly improved anxiety-like behavior on the OFT and EPM; treatment with chemical psilocybin at the same dose was not as effective. In Study 2 of the current project, psilocybin 4.4 mg/kg was not effective in improving anxiety-like behavior in SAPAP3-KO mice. The effect of psychedelic mushroom extract remains to be studied in juvenile SAPAP3-KO mice.

The MBT showed intriguing and highly significant effects of genotype in both sexes. In Study 1 and Study 2, juvenile SAPAP3-KO mice homozygous for the deletion, buried significantly fewer marbles over 30 minutes than WT mice. Similar findings were reported by (Brownstien et al., 2024) in adult SAPAP3-KO mice. Digging, burrowing, and burying are natural behaviors in rodents and not necessarily indicative of pathology (Jirkof, 2014). Nevertheless the MBT is widely used as a predictive test for the potential efficacy of compounds in treating OCD (Matsushima et al., 2009; Odland et al., 2021) and as a measure of anxiolytic effects (Njung’e and Handley, 1991). However, the relevance of the test has been questioned on the grounds of face and predictive validity (Jimenez-Gomez et al., 2011; De Brouwer et al., 2019). In the current study, reduced marble burying in SAPA2-KO mice was related to a lower level of activity and a higher level of anxiety as indicated by significant correlations number of marbles buried with activity on the OFT and time spent in the open arms of the EPM respectively.

In adult SAPAP-3 KO mice treated with a single injection of psilocybin (4.4 mg/kg) or psychedelic mushroom extract containing the same dose of psilocybin, marble burying was significantly increased by the treatment (Brownstien et al., 2024). The MBT was performed by (Brownstien et al., 2024) 3 days after the administration of the mushroom extract. At this time, the pathological self-grooming and head-body twitches characteristic of adult HOM SAPAP3-KO mice were also alleviated by psychedelic mushroom extract and by psilocybin. In contrast to the effect of psilocybin on marble burying in adult SAPAP3-KO mice, in Study 2 of the current project, there was no effect of psilocybin on the same phenotype in juvenile mice. This suggests that the effect of psilocybin on behavioral manifestations of SAPAP3 deletion may be related to the maturation of brain systems. Brain maturation, especially in circuits implicated in compulsive behaviors or OCD-like symptoms, may influence psilocybin’s efficacy. Psilocybin interacts with serotonin receptors which are known to play a critical role in mood and behavior. During maturation, the distribution, density, and sensitivity of these receptors undergo significant changes, potentially affecting how psilocybin modulates neural activity (Murrin et al., 2007). In adult SAPAP3-KO mice, mature receptor networks and more developed neural circuits may provide a suitable substrate for psilocybin’s action, particularly in areas like the prefrontal cortex or amygdala, which are crucial in regulating compulsive behavior. The synaptic plasticity and connectivity patterns in a mature brain could allow psilocybin to exert more profound effects on neural circuit function, which might be less developed or stable in juvenile mice.

The results of the tube dominance test in Study 1 revealed greater assertiveness of male HOM mice than male WT mice but no significant effect in females. In Study 2, male and female HOM mice were both more assertive than WT mice. Greater assertiveness of HOM mice on the tube dominance test is unexpected in view of the reported association of SAPAP3 deletion with lower sociability (Rajkumar, 2023), although it should be noted that in Study 1 of the current project, genotype-associated impairment of social interaction was not observed in SAPAP3-KO mice. A relationship of winning on the tube dominance test to activity on the OFT and anxiety level on the EPM is suggested by the results of Study 2 but not those of Study 1, so that this relationship remains an open question.

Buried food seeking is a frequently used test in olfaction research in mice (Yang and Crawley, 2009; Machado et al., 2018). The Buried Oreo test used in our studies combines the predilection of mice to unearth buried food substances with the preference of mice for sweet taste. This preference is a basis for the sucrose preference test that is widely used as a measure of anhedonia in mice (Markov, 2022). In both studies reported here, HOM juvenile SAPAP3-KO mice of both sexes unearthed significantly fewer buried Oreo cookies than WT mice. There was no difference between genotypes in either study in number of unearthed Oreos that were eaten although there was a trend in this direction in Study 2. Thus, it is not entirely clear whether the lower tendency to unearth buried Oreos in the HOM mice is a manifestation of anhedonia or lack of motivation, Psilocybin treatment in Study 2 did not alter number of Oreos unearthed or eaten.

Contrary to these positive findings, there was no effect of SAPAP3-KO genotype on the FST reflecting absence of depressive-like features (Study 1), on social exploration (Study 1) or on two cognitive measures, the Novel Object Recognition test (Study 1) and Y-Maze (Studies 1 and 2). Lack of difference on the FST, which is sensitive to activity, supports the finding that in other tests (such as EPM and MBT) the significant effect of genotype was not a consequence of reduced activity in HOM SAPAP3-KO mice.

Our findings reveal significant effects of genotype on synaptic plasticity markers in adult SAPAP3-KO mice in the frontal cortex, amygdala, hippocampus, and striatum. In post-hoc analyses by sex, the synaptic proteins showing significantly higher levels in male HOM mice were GAP43 (in frontal cortex and amygdala), synaptophysin (in frontal cortex), and SV2A (in hippocampus, amygdala, and striatum). In female HOM mice, SV2A was significantly elevated in the frontal cortex compared to female WT mice. Although there was a significant effect of genotype on PSD95 in the frontal cortex, levels of this synaptic protein were not significantly different by post-hoc testing in either sex separately in any of the four brain areas. No significant effects of genotype or sex on synaptic protein levels were observed in juvenile SAPAP3 KO mice aged 12 weeks. Our findings regarding genotype are partially in accordance with those of (Glorie et al., 2020), who used positron emission tomography (PET) to examine SV2A levels in the cortex, hippocampus, thalamus, and striatum of female SAPAP3-KO and WT mice. (Glorie et al., 2020) found that at the age of 3 months, the female SAPAP3-KO mice they studied manifested significantly lower SV2A binding compared to WT littermates in the cortex and hippocampus. thalamus and striatum. Aging in WT mice was associated with a significant (p < 0.001) decline in SV2A binding throughout the brain, whereas in SAPAP3 KO mice, this decline was confined to the corticostriatal level (Glorie et al., 2020).

The increase in synaptic proteins in SAPAP3-KO mice points to enhanced synaptic growth and vesicle-associated plasticity in the affected regions. GAP43 is a marker of axonal growth and synaptic remodeling, while synaptophysin and SV2A are essential components of synaptic vesicle regulation and neurotransmitter release (Holahan, 2015; Rossi et al., 2022; Bera et al., 2023). SAPAP3 is a scaffold protein that interacts directly with PSD95, as its name - SAP90/PSD95-associated protein 3 - suggests. Importantly, SAPAP3 is a post-synaptic protein located in the post-synaptic density and is highly expressed in the striatum. Despite this, our findings showed less striking changes in PSD95 levels in male or female adult KO mice. This suggests that the alterations we observed may be secondary to disrupted signaling pathways involving the cortex-thalamus-striatum circuit. The increased synaptic proteins in the cortex, hippocampus, and amygdala of male SAPAP3-KO mice could reflect compensatory changes in response to dysregulated inputs from the striatum (Bai et al., 2022). Thus, plasticity alterations in SAPAP3-KO mice may be presynaptic, affecting synaptic transmission and connectivity rather than postsynaptic structures. This is intriguing given that synaptic pruning, a normal developmental process that refines neural circuits during adolescence, involves the selective removal of unnecessary synapses, typically reducing plasticity in adult brain regions (Kirkland et al., 2024). In this context, it is of interest that (Glorie et al., 2020) found a decrease in SV2A binding in adult compared to juvenile SAPAP3-WT mice. This decrease was less consistent in SAPAP3-KO mice.

As noted, the observed increases in synaptic proteins occurred only in adult SAPAP3-KO mice. The fact that these changes were absent in younger mice supports the idea of an age-dependent progression of pathology, potentially related to abnormal synaptic pruning. In healthy humans, synaptic pruning during adolescence serves to eliminate excess synapses, optimizing neural networks for efficient processing (Kirkland et al., 2024). Disruptions in this process have been implicated in various neuropsychiatric disorders (Eltokhi et al., 2020; Westacott and Wilkinson, 2022). Specifically, delayed or abnormal pruning has been associated with atypical neural development, as seen in humans with higher neuropsychopathological (NP) factor scores, where a reduction in gray matter volume is inhibited during adolescence (Xie et al., 2023). These findings suggest that similar processes may occur in male SAPAP3-KO mice, with altered synaptic pruning contributing to the pathological increase in synaptic proteins in adulthood. The increase in plasticity markers in older SAPAP3-KO mice might reflect a compensatory or maladaptive response to impaired pruning, particularly in regions involved in emotional and cognitive regulation. This may explain why the obsessive grooming behaviors, a hallmark of this model, emerge in adulthood rather than earlier in life.

Despite molecular differences, male and female mice showed similar OCD-like behaviors (Brownstien et al., 2024), indicating that behavioral phenotypes were not strongly influenced by sex. The less striking synaptic protein increases in females, despite comparable behaviors, may be due to sex-specific neurodevelopmental processes. Studies show distinct patterns of synaptic pruning and plasticity between sexes, influenced by hormones like testosterone and estrogen, which shape brain plasticity differently (Han et al., 2021). Males undergo more robust synaptic pruning in regions like the nucleus accumbens during adolescence (Kirkland et al., 2024), possibly explaining the observed increase in plasticity markers in older male mice.

## Conclusions

Our findings demonstrate, in two separate mouse cohorts, high levels of anxiety-like behavior in juvenile SAPAP3-KO mice that are homozygous for deletion of the gene, as well as other behavioral differences. The high levels of anxiety that we observed were not responsive to treatment with psilocybin. The excessive self-grooming observed in adult SAPAP3-KO mice is regarded as possibly modeling OCD; our current findings are analogous to the prodromal anxiety observed in patients with OCD and can provide an important basis for further studies. Our intriguing findings regarding increased neuroplasticity in adult, but not juvenile, male SAPAP3- mice are a basis for further research to elucidate the mechanisms underlying these changes and their implications for OCD.

## Funding

The Hadassah BrainLabs Center for Psychedelic Research was founded with the support of Negev Labs. This work was supported in part by Back of the Yards algae sciences (BYAS) and Parow Entheobiosciences (ParowBio).

## Acknowledgments

Prof. Tamir Ben-Hur provided valuable advice on synaptic protein analyses, Masha Chaykin expertly assisted with Western Blot assays. Bar Orian assisted with the behavioral assays.

## Conflict of Interest

BL is a consultant to Back of the Yards algae sciences (BYAS) and Parow Entheobiosciences (ParowBio). ParowBio has submitted a patent application (WO 2023\164092 A1) on the use of psilocybin-containing psychedelic mushroom extract to treat psychiatric disorders including OCD on which BL is a co-inventor. None of the other authors have any conflict of interest.

## Data Availability Statement

Data is available to qualified investigators on request from the corresponding authors.

## Notes

### Summary of Updates

Significant clarifications of behavioral and statistical methodology have been provided. Representative immunoblot images have been added. Supplemental files have been updated

