## SUPPLEMENTARY INFORMATION for "Premorbid Characteristics of the SAPAP3-Mouse Model of Obsessive-Compulsive Disorder: Behavior, Neuroplasticity, and Psilocybin Treatment"

**Lazar et al:**

**SUPPLEMENTARY INFORMATION**

**1. Behavioral Methods**

Behavioral methods are given in the order in which the results are presented in the Results Section.

- **Open Field Test (OFT)**

The OFT was performed immediately after the MBT to evaluate locomotor activity. The apparatus consisted of a square wooden arena (50 × 50 × 40 cm) with white walls and floor. Mice were placed individually in the center of the open field and allowed to freely explore the apparatus for 10 min. A camera was used to monitor movement. The total distance traveled (millimeters) was measured by the Ethovision XT-12 Video Tracking System (Noldus Information Technology BV). The ﬂoor space was divided into peripheral and central regions by the system and the time spent in each area was recorded. After each test, the arena was cleaned with a 70% alcohol solution (Lok et al., 2013; Lotan et al., 2018).

- **Elevated Plus Maze Test (EPM)**

Anxiety-like behavior was measured using the EPM. The apparatus used is in the configuration of a + and comprises two open arms (25 x 5 x 0.5 cm) across from each other and perpendicular to two closed arms (25 x 5 x 16 cm) with a center platform (5 x 5 x 0.5 cm). The open arms have a very small (0.5 cm) wall to decrease the number of falls, whereas the closed arms have a high (16 cm) wall to enclose the arm. The entire apparatus is 75 cm above the floor. The animal is placed on the central platform and is allowed to explore the arena freely for 6 minutes. The EthoVision XT-12 Video Tracking System (Noldus Information Technology BV) tracks the animals and measures the duration and frequency of their presence in the closed and open arms. Before starting a new EPM test the arena was cleaned with a 70% alcohol solution (Komada et al., 2008; Lifschytz et al., 2012).

- **Marble Burying Test (MBT)**

The MBT was performed in transparent cages containing ~4.5 cm fine sawdust. Twenty glass marbles were placed equidistant from each other in a 5 × 4 pattern. The experiment was done under dim light in a quiet room to reduce the influence of anxiety on behavior. The mice were left in the cage with the marbles and after a 30-min period, the buried marbles were counted. A marble was considered buried if two-thirds or more of its size was covered with sawdust. The outcome measure in this test is the number of marbles buried in 30 minutes (Singh et al., 2023).

- **Tube Dominance Test**

The apparatus for the Tube Dominance Test consists of a 30 cm transparent tube with a 3.5 cm internal diameter. Before the test each animal underwent 2 days of training trials, in which they explored the entire tube length (without the presence of an opponent) and entered and emerged from the tube 5 times. On the day of the test at the beginning of each trial, the test animal and another cage mate (stimulus animal) were introduced simultaneously into the tube, each from a different edge of the tube. Altogether, there were at least 3 trials for each test animal. A trial will be completed when one of the animals (test and stimulus) is defeated and has been pushed back until two of the legs are outside the tube. Otherwise, if there was no defeat the trial would end after 2 minutes. During each of the trials, a test animal would receive a grade of 0 if defeated, 0.5 if draw and 1 if won. Each mouse's score is determined by the percentage of wins, which is calculated by dividing the total points earned across all trials by the total number of possible wins (Zhou et al., 2017; Fan et al., 2019).

- **Buried Oreo Test**

Anhedonia-related behavior was measured using the Buried Oreo Test. In this test, for habituation, one half of an Oreo cookie for every 3 mice was inserted into the mouse’s home cage (not buried) for 2 hours. Home cages needed to be replaced after habituation to remove any odor of the cookie from the mice’s environment. On the next day the mouse was placed in a different cage with 6-8 cm height sawdust, alone, for 30 min of habituation. At the end of the habituation the mouse was removed from the cage and half of an oreo cookie was buried approximately 2 cm under the bedding. The mouse was then put back into the cage for a 10 min test. The test ends when the mouse eats the cookie or when 10 min passed, whatever comes first. The test is manually scored. For digging up the cookie and finding it, the mouse receives 1 point, if he did not dig it up, 0 points. For eating the cookie, the mouse receives 1 point, if he does not eat it, 0 points (Yang and Crawley, 2009; Machado et al., 2018).

- **Forced Swim Test (FST)**

In the FST, the mouse is placed in a water-filled, transparent, round plexiglass tank. For the duration of the 6-minute test period, the time spent by the animal in activity (active swimming) versus immobility (passive floating) is tracked and recorded by the the EthoVisionXT-12 video-imaging system (Noldus Information Technology BV)., as well as the frequency and duration of activity/ inactivity bouts. These variables serve as the outcome measure of the test(Lifschytz et al., 2006).

- **Y Maze Test**

Working memory and exploratory activity were measured using a Y-maze apparatus. The apparatus consisted of a Y shaped arena (arm length: 40 cm, arm bottom width: 3 cm, arm upper width: 13 cm, height of wall: 15 cm). Each mouse was placed in the central area. The number of entries into the arms and alterations were recorded for 6 min with the EthoVisionXT-12 video-imaging system (Noldus Information Technology BV). Working memory was calculated as the number of correct alternations/total number of alternations (Yoshizaki et al., 2020).

- **Novel Object Recognition Test (NOR)**

NOR, by evaluating differences in the exploration time of novel and familiar objects, provides a measure for recognition and episodic memory. The mouse is placed inside a transparent plexiglass box 50cm in all dimensions, with two similar “familiar” objects for a habituation stage of 10 minutes in opposite. In the test stage, the animal is re-entered to the arena for a 4-minute session, now with one “familiar” and one “different “novel” object. The duration of time spent near each object and the frequency and number of approaches are tracked by the the EthoVisionXT-12 video-imaging system (Noldus Information Technology BV). and serve as outcome measures of the test (Lotan et al., 2018).

- **Social Exploration Test**

This test evaluates how sociable mouse is when it meets a new mouse for the first time. First, the mouse has a habituation time where’s it spends 10 minutes alone in a new cage to get used to it. Then, the mouse meets with a new mouse, from the same gender, and the two mice are put together in the same cage for another 2 minutes.

The test tracks the total percentage of time the mouse in study is spend interacting (sniffing) with the new mouse.

The tracking is done with EthoVisionXT-12 video-imaging system (Noldus Information Technology BV), and a new litter is change between mice.

**2. Supplementary Tables**

**Supplementary Table 1:** Baseline features of mice participating in the experiments

**Study 1**

|  | **Female** | | | **Male** | | |
| --- | --- | --- | --- | --- | --- | --- |
| **Genotype** | **HOM** | **HET** | **WT** | **HOM** | **HET** | **WT** |
| **N** | 24 | 21 | 29 | 20 | 21 | 26 |
| **Age (Weeks±SD)** | 10.94 ± 0.9 | 10.82 ± 0.95 | 10.81 ± 1.01 | 10.96 ± 0.87 | 10.86 ± 0.8 | 10.9 ± 0.78 |
| **Weight (gm±SD)** | 19 ± 2.3 | 18.54 ± 2.2 | 16.97 ± 3.9 | 23.25 ± 2.9 | 23.29 ± 5.6 | 25.73 ± 1.64 |

No differences were statistically significant

**Study 2**

|  | **Female** | | **Male** | |
| --- | --- | --- | --- | --- |
| **Genotype** | **HOM** | **WT** | **HOM** | **WT** |
| **N** | 16 | 16 | 16 | 16 |
| **Age (Weeks±SD)** | 12.06±085 | 11.5±0.82 | 12.05±0.85 | 11.79±0.88 |
| **Weight (gm±SD)** | 18.36 ±1.08 | 21.2 ±1.58 | 24.83 ±2.3 | 27.2 ±1.79 |
| **Treatment** |  |  |  |  |
| **Vehicle (N)** | 16 | 16 | 16 | 16 |
| **Psilocybin (N)** | 16 | 16 | 16 | 16 |

No differences were statistically significant

**3. Supplementary Figures**

**Supplementary Fig. 1**

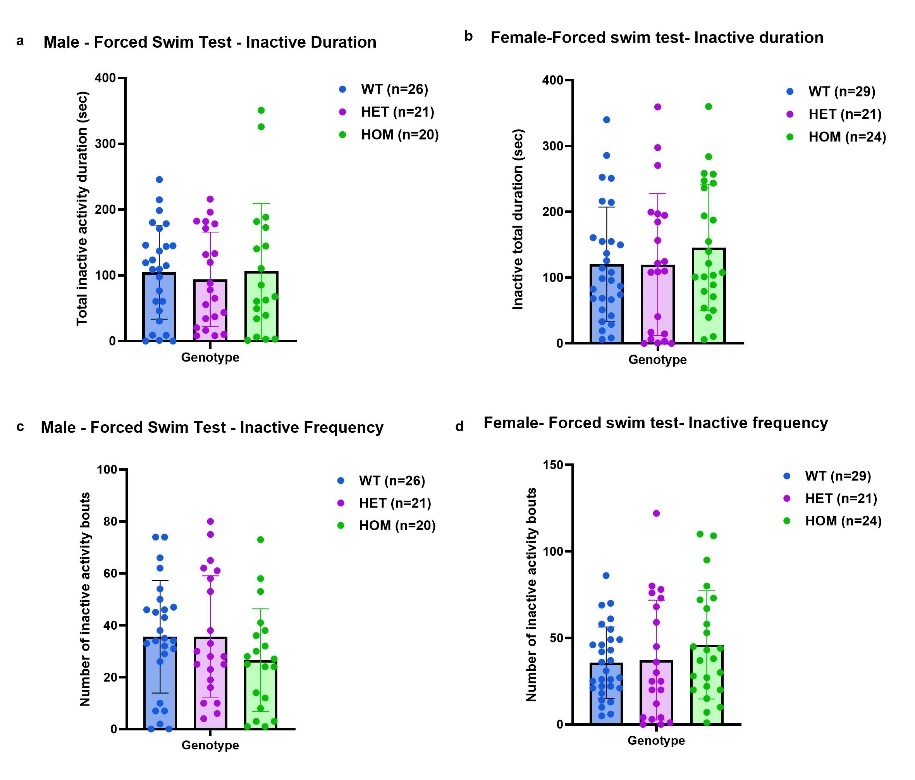

**Supplementary Fig. 1:** Study 1: Forced swim test comparing WT, HET, and HOM SAPAP3-KO mice. Inactive duration in males (a) and females (b), and inactive frequency in males (c) and females (d). No significant genotype or gender effects

**Supplementary Fig. 2**

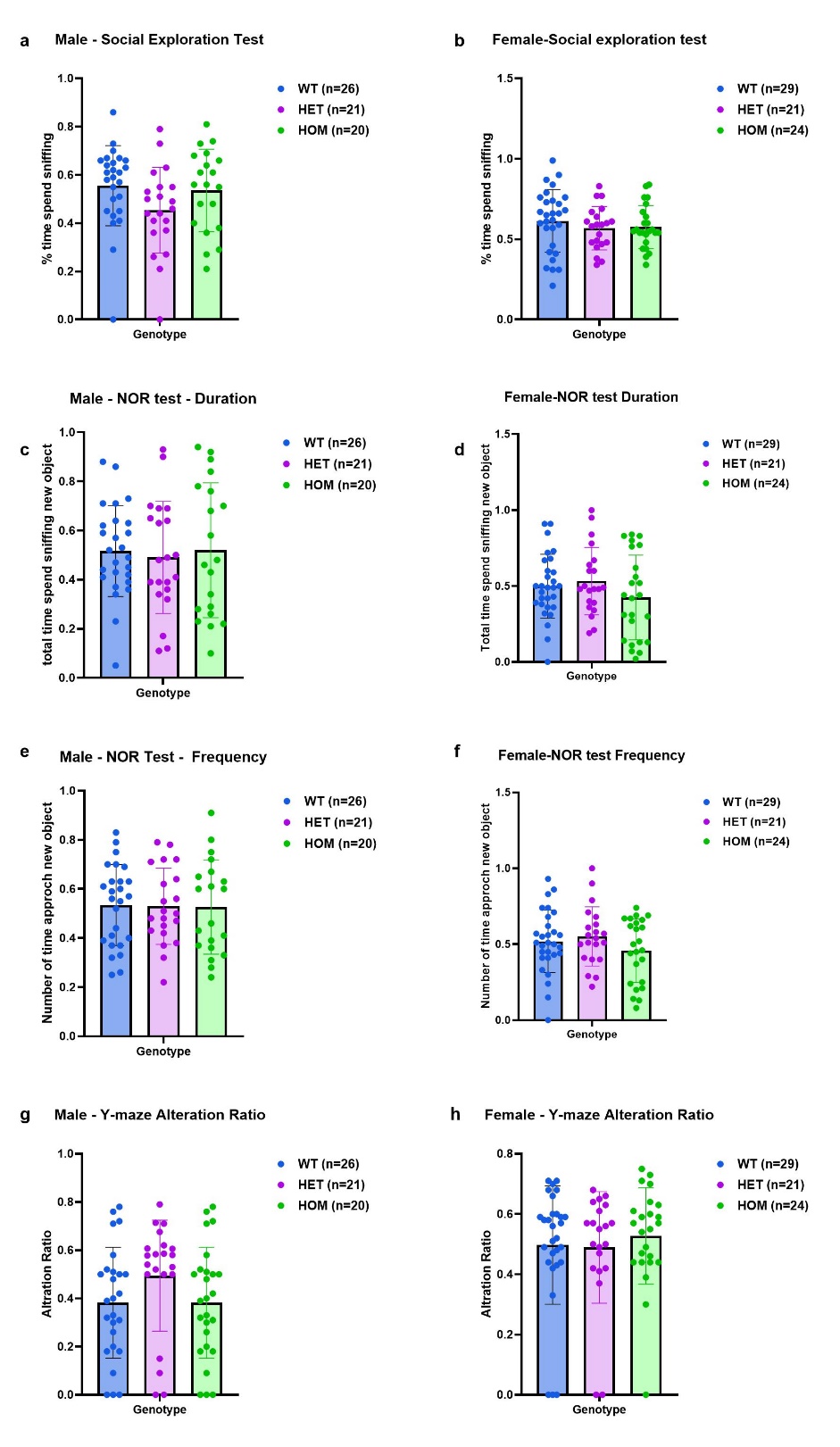

**Supplementary Fig. 2:** Study 1: Social exploration test (a, b), NOR test duration (c, d), NOR test frequency (e, f) Y-maze alternation ratio. No significant genotype or gender effects

**Supplementary Fig. 3**

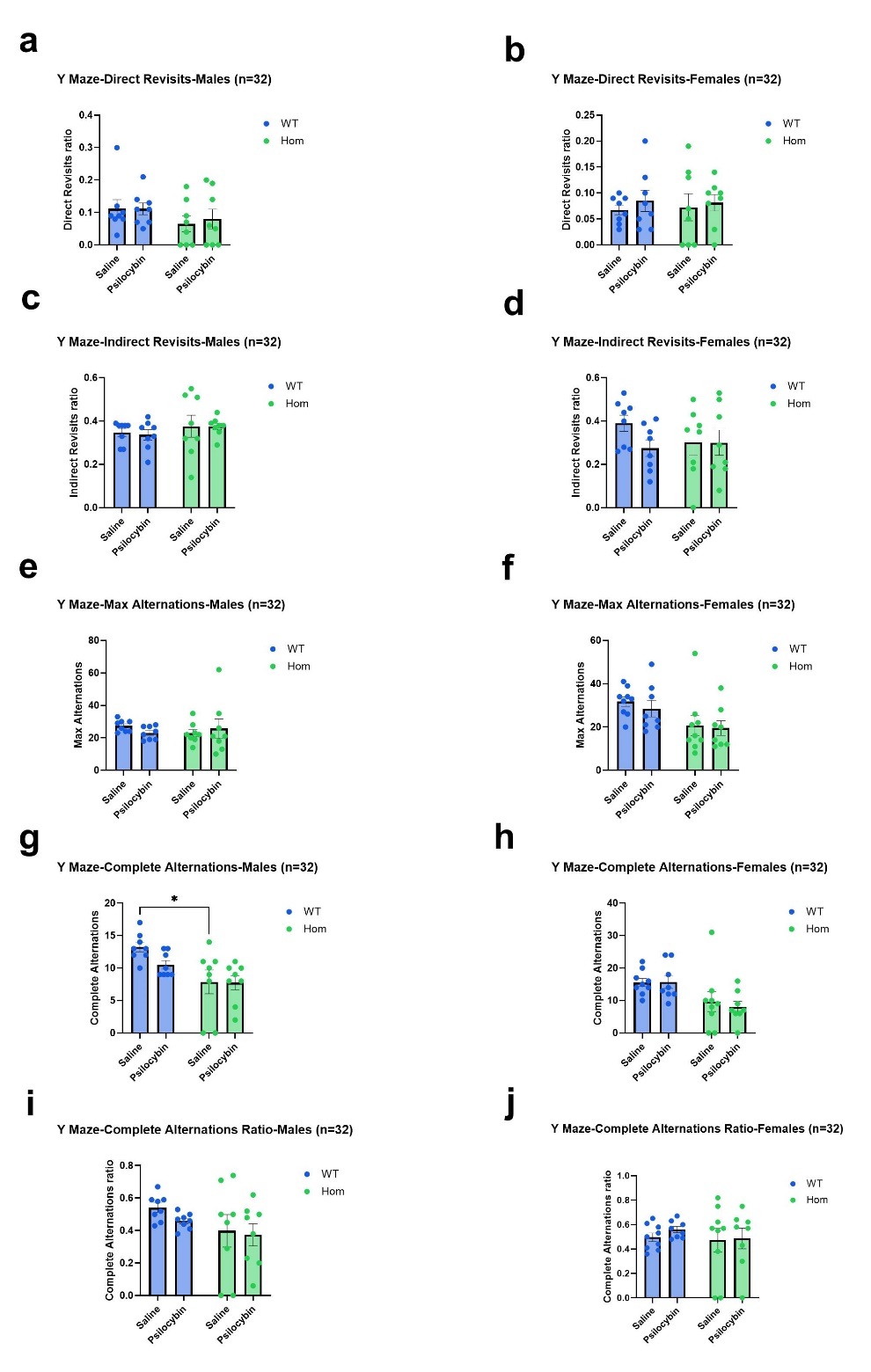

**Supplementary Fig. 3:** Study 2: Y Maze - Direct revisits in males (a) and females (b), Indirect revisits in males (c) and females (d), Max alternations in males (e) and females (f), Complete alternations in males (g) and females (h) and complete alternation ratio in males (i) and females (j), in WT and SAPAP3-KO mice, under treatment with saline or psilocybin.

e,f - Significant effect of genotype on max alternations in females (F=7.77; df 1,28; p=0.009) but not in males.

g, h - Complete alternations in a) males (F=9.91; df 1,28; p=0.0039) and b) females (F=26.51; df 1,28; p<0.0001; **p<0.01, Saline WT vs. Saline HOM; ^+^p<0.05, Psilocybin WT vs. Psilocybin HOM).

i, j - No significant differences in alternation ratio in a) males or b) females

**Supplementary Fig. 4**

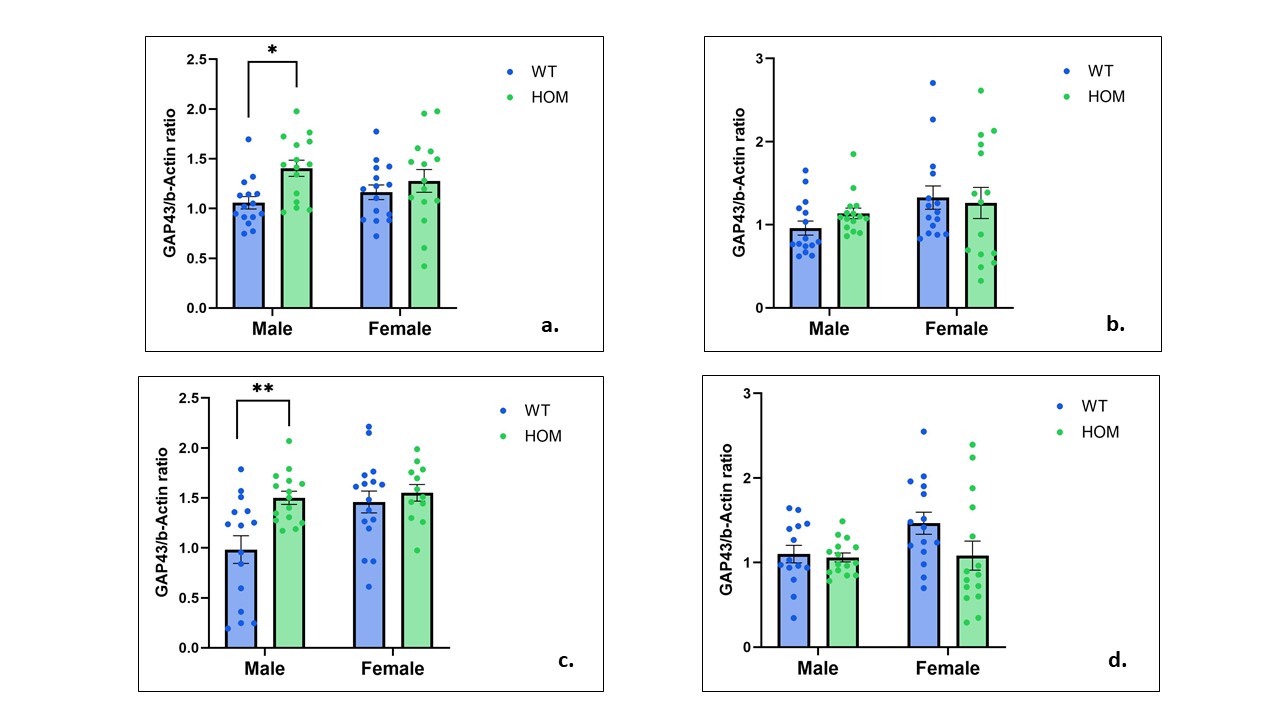

**Supplementary Fig. 4:** Study 1: Two-way ANOVA of GAP43 levels in the frontal cortex (a), hippocampus (b), amygdala (c) and striatum (d). Significant effects of genotype observed for GAP43 in the frontal cortex (F=7.30, df 1,56, p=0.009) and amygdala (F=8.24, df 1,54, p=0.005), post-hoc significance for male HOM vs. male WT was 0.02 and 0.04 for each area respectively.

Significant sex effects on GAP43 in the amygdala (F=6.15, df 1,54, p=0.01).

**Supplementary Fig. 5**

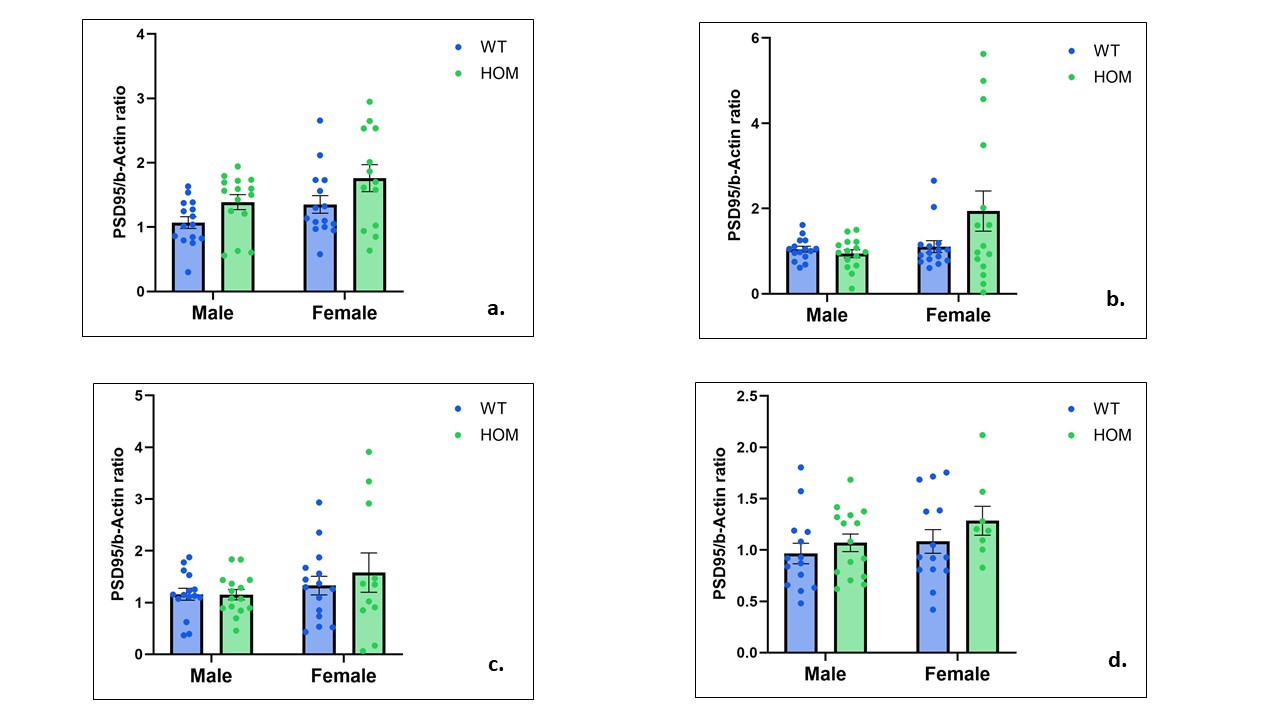

**Supplementary Fig. 5:** Study 1: Two-way ANOVA of PSD95 levels in the frontal cortex (a), hippocampus (b), amygdala (c) and striatum (d). Significant effect of genotype in the frontal cortex (F=6.70, df 1,54, p=0.01). Post-hoc tests were not significant in either sex separately.

Significant sex effects on PSD95 in the frontal cortex (F=5.43, df 1,54, p=0.02) and hippocampus (F=4.38, df 1,56, p=0.04)

**Supplementary Fig. 6**
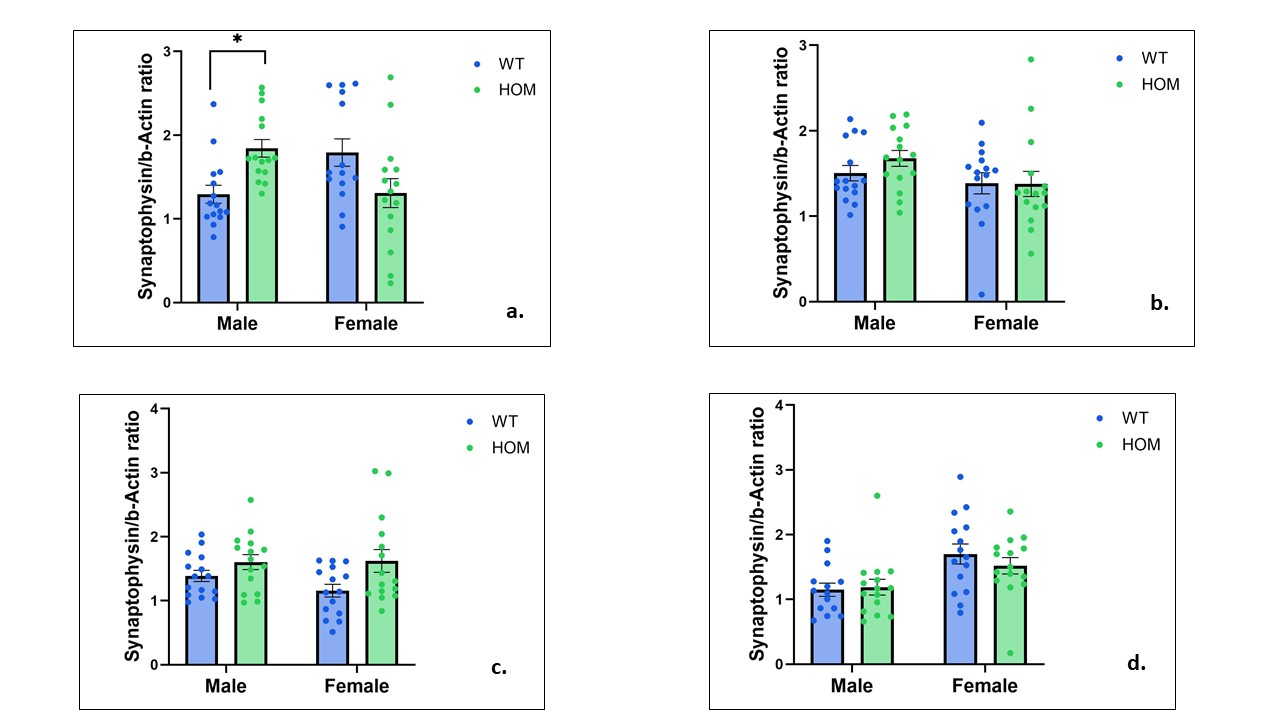

**Supplementary Fig. 6:** Study 1: Two-way ANOVA of synaptophysin levels in the frontal cortex (a), hippocampus (b), amygdala (c) and striatum (d). Significant effect of genotype for synaptophysin in the amygdala (F=7.34, d 1,56, p=0.008). Post-hoc tests not significant in either gender separately.

Significant sex effect on synaptophysin in the striatum (F=11.77, df 1,55, p=0.001).

Significant sex x genotype interaction for synaptophysin the frontal cortex (F=13.51 df 1,55,

**Supplementary Fig. 7**
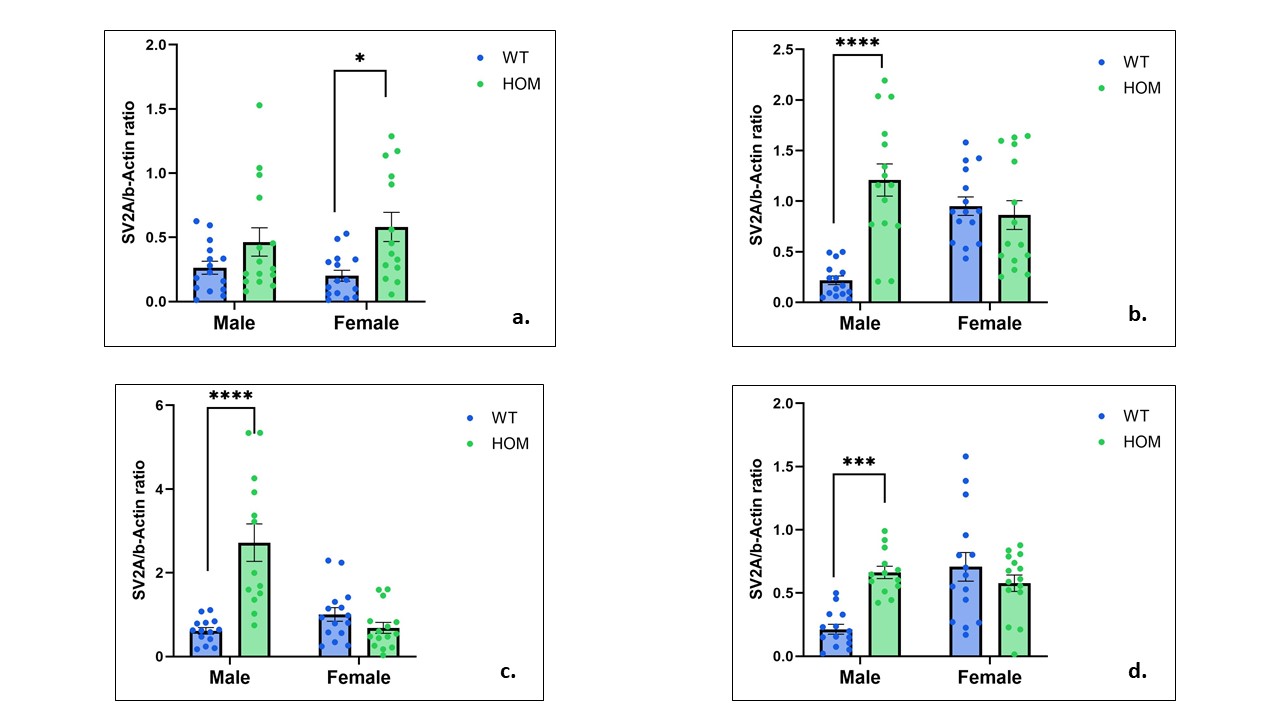

**Supplementary Fig. 7:** Study 1: Two-way ANOVA of SV2A levels in the frontal cortex (a), hippocampus (b), amygdala (c) and striatum (d). Significant effect of genotype on SV2A levels in the frontal cortex (F=17.22, df 1,55, p=0.001), (Female HOM mice manifest significantly higher SV2A levels than female WT mice, Tukey post-hoc, *p<0.05). Genotype effects also highly significant in the hippocampus (F=14.67, df 1,56, p=0.0003), amygdala (F=14.48, df 1,53, p=0.0004) and striatum (F=4.59, df 1,53, p=0.03). Post-hoc tests showed that SV2A levels were significantly higher in male HOM than male WT mice in all three areas (***p<0.001, ****p<0.0001).

Significant sex effects on SV2A levels in the amygdala (F=12.32, df 1,53, p=0.0009) and striatum (F=7.48, df 1,53, p=0.008).

Significant sex x genotype interaction for SV2A in the hippocampus (F=20,94, df 1,56, p=,0.0001), amygdala (F=26.75, df 1,53, p<0.0001) and striatum (F=15.08, df 1,53, p=0.0003)

**Supplementary Fig. 8.**

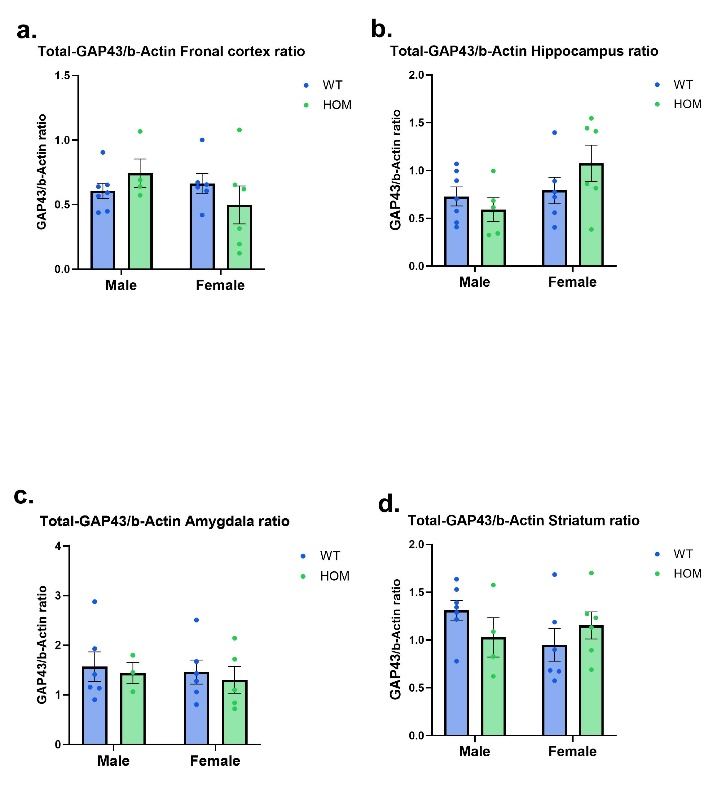

**Supplementary Fig. 8:** Study 2: Two-way ANOVA of GAP43 levels in the frontal cortex (a), hippocampus (b), amygdala (c) and striatum (d). No statistically significant effects

**Supplementary Fig. 9.**

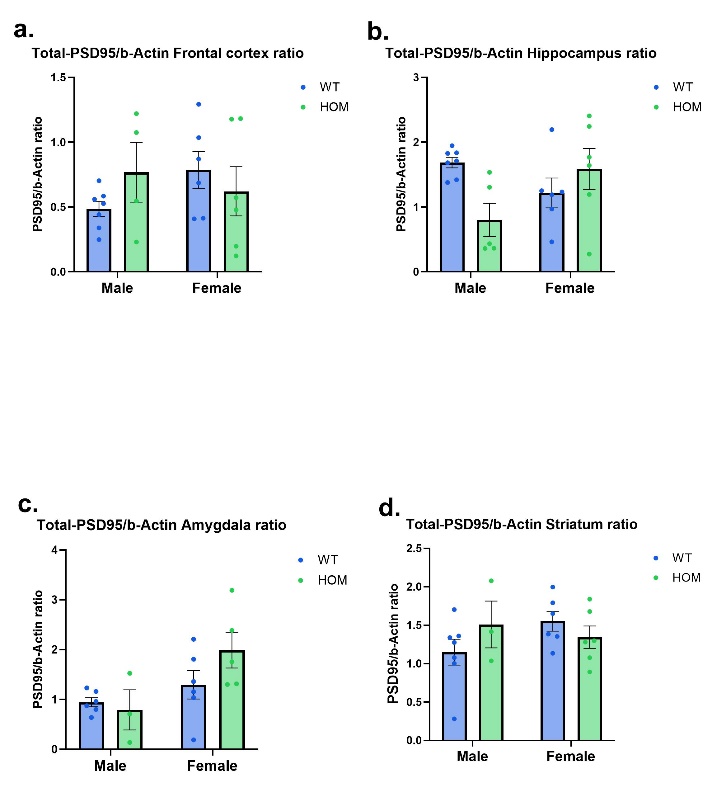

**Supplementary Fig. 9:** Study 2. Two-way ANOVA of PSD95 levels in the frontal cortex (a), hippocampus (b), amygdala (c) and striatum (d). Significant gender effect for PSD95 in the amygdala (F=7.10, df 1,16, p=0.01) and a significant gender by genotype interaction for PSD95 in the hippocampus (F=7.54, df 1,20, p=0.01). No post-hoc tests were statistically significant.

**Supplementary Fig. 10.**

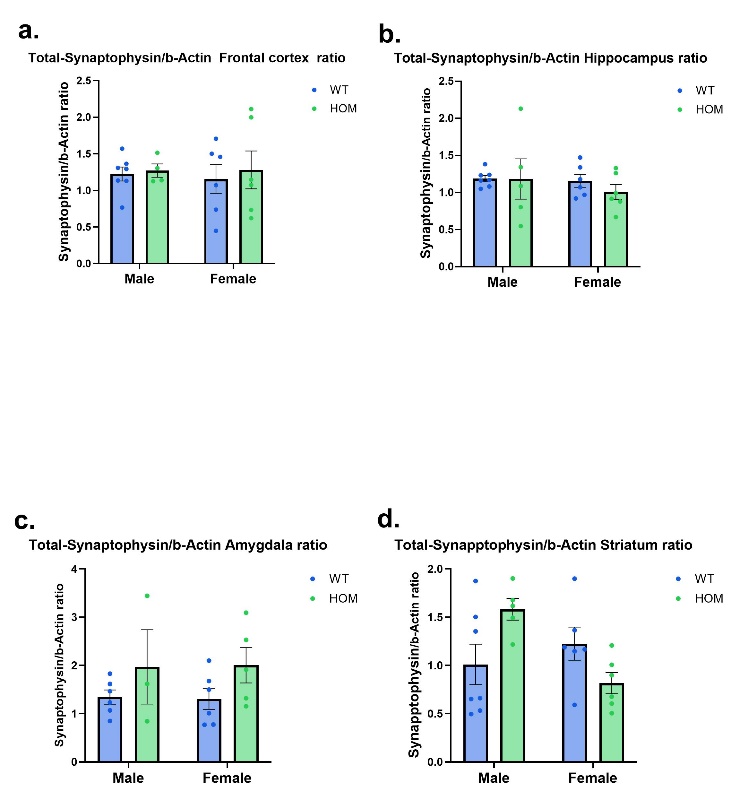

**Supplementary Fig. 10:** Study 2. Two-way ANOVA of synaptophysin levels in the frontal cortex (a), hippocampus (b), amygdala (c) and striatum (d). Significant gender by genotype interaction for synaptophysin in the striatum (F=7.49, df 1,20, p=0.008). No post-hoc tests were statistically significant.

**Supplementary Fig. 11**

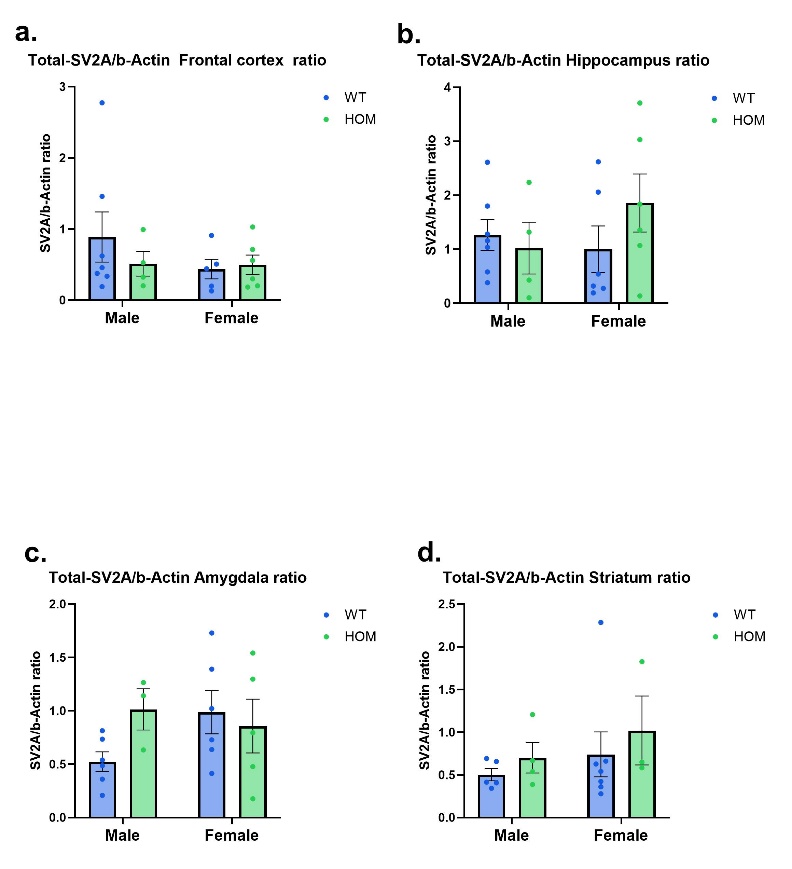

**Supplementary Fig. 11:** Study 2. Two-way ANOVA of SV2A levels in the frontal cortex (a), hippocampus (b), amygdala (c) and striatum (d). No comparisons were statistically significant.

**Supplementary Fig. 12.**

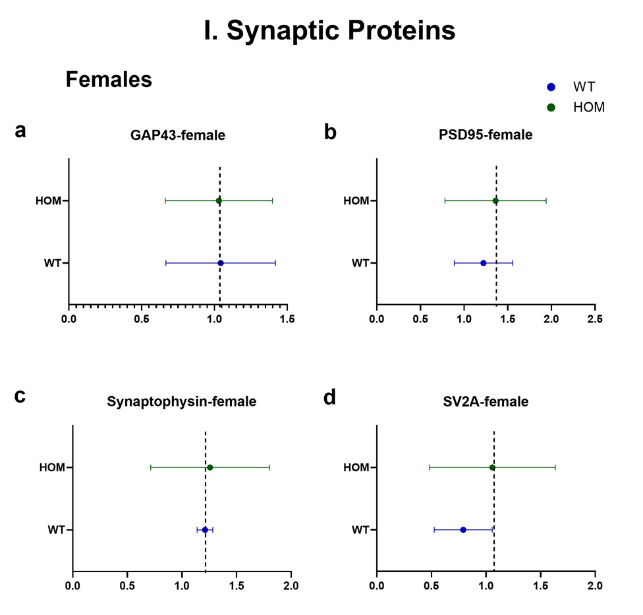

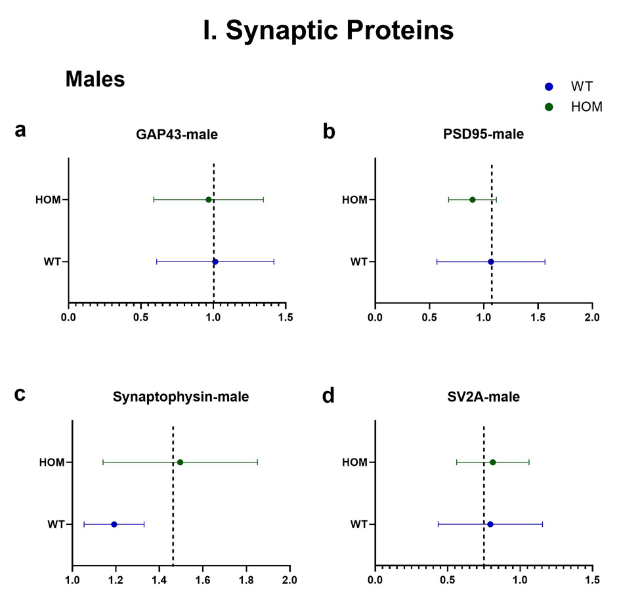

**Supplementary Fig. 12:** Study 2. Nested analysis of synaptic proteins (GAP43, PSD95, synaptophysin, and SV2A) across 4 brain areas (frontal cortex, hippocampus, amygdala, striatum) in male (left) and female (right) HOM vs. WT SAPAP3-KO mice.

II. Nested analysis of 4 synaptic proteins (GAP43, PSD95, synaptophysin, and SV2A) within each of 4 brain areas (frontal cortex (a), hippocampus (b), amygdala (c), striatum (d)) in male and female HOM vs. WT SAPAP3-KO mice. Nested t-test did not show significant differences

**Supplementary Fig. 13.**

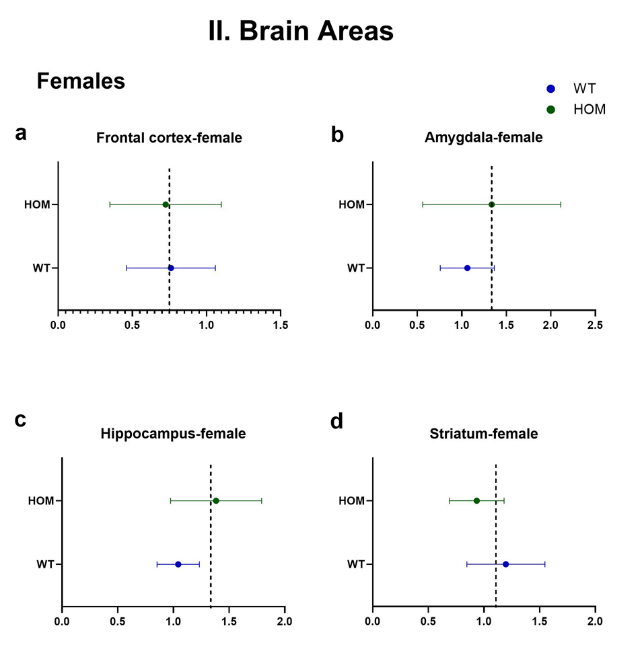

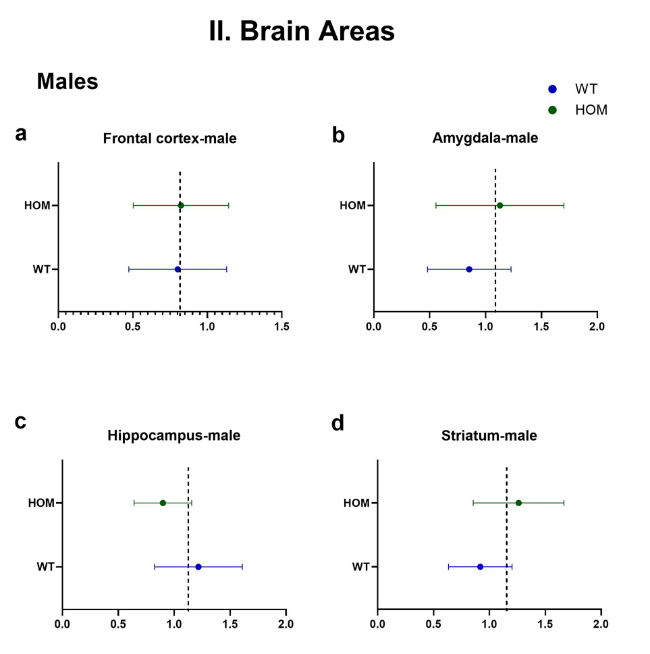

**Supplementary Fig. 13:** Study 2. Nested analysis of synaptic proteins (GAP43, PSD95, synaptophysin, and SV2A) within each of 4 brain areas (frontal cortex, hippocampus, amygdala, striatum) separately in male (left) and female (right) HOM vs. WT SAPAP3-KO mice. Nested t-test did not show significant differences

**Supplementary Fig. 14A**

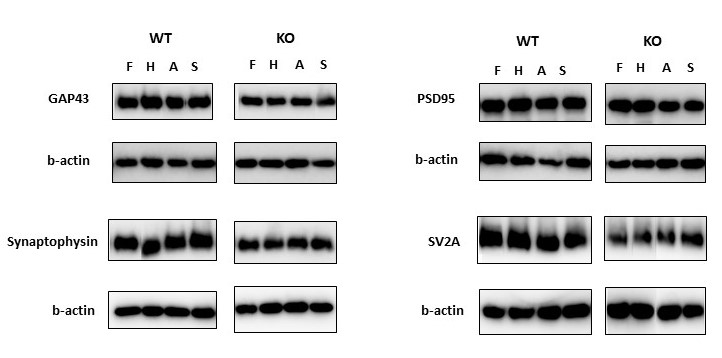

**Supplementary Fig. 14A:** Representative immunoblots of GAP43, PSD95, Synaptophysin, SV2A and b-actin protein expression across 4 brain areas (frontal cortex (F), hippocampus (H), amygdala (A), and striatum (S)) by genotype (SAPAP3-KO and WT) in adult male mice.

**Supplementary Fig. 14B**

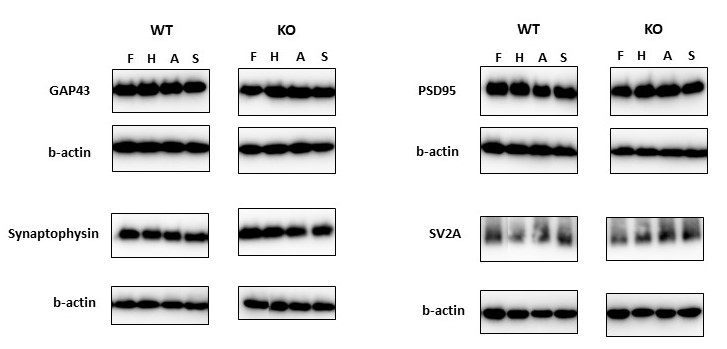

**Supplementary Fig. 14B:** Representative immunoblots of GAP43, PSD95, Synaptophysin, SV2A and b-actin protein expression across 4 brain areas (frontal cortex (F), hippocampus (H), amygdala (A), and striatum (S)) by genotype (SAPAP3-KO and WT) in adult female mice.

**Supplementary Fig. 14C**

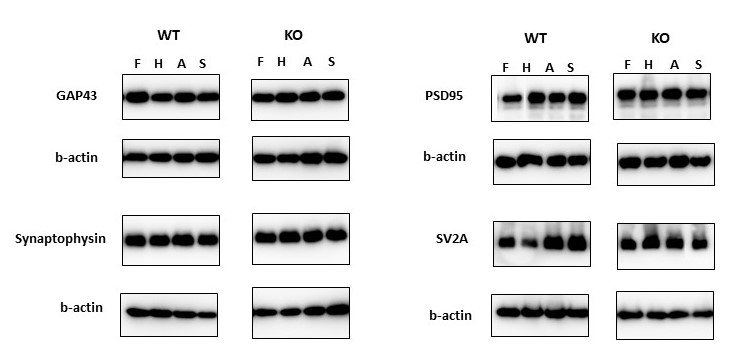

**Supplementary Fig. 14C:** Representative immunoblots of GAP43, PSD95, Synaptophysin, SV2A and b-actin protein expression across 4 brain areas (frontal cortex (F), hippocampus (H), amygdala (A), and striatum (S)) by genotype (SAPAP3-KO and WT) in juvenile male mice.

**Supplementary Fig. 14D**

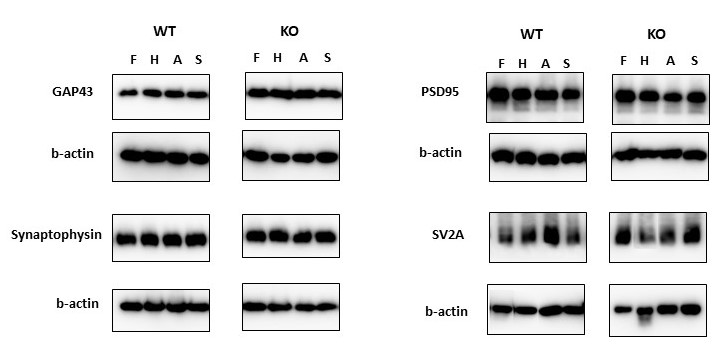

**Supplementary Fig. 14D:** Representative immunoblots of GAP43, PSD95, Synaptophysin, SV2A and b-actin protein expression across 4 brain areas (frontal cortex (F), hippocampus (H), amygdala (A), and striatum (S)) by genotype (SAPAP3-KO and WT) in juvenile female mice.
